# Selective Enhancement of the Interneuron Network and Gamma-Band Power via GluN2C/GluN2D NMDA Receptor Potentiation

**DOI:** 10.1101/2024.11.05.622179

**Authors:** Chad R. Camp, Tue G. Banke, Hao Xing, Kuai Yu, Riley E. Perszyk, Matthew P. Epplin, Nicholas S. Akins, Jing Zhang, Tim A. Benke, Hongjie Yuan, Dennis C. Liotta, Stephen F. Traynelis

## Abstract

N-methyl-D-aspartate receptors (NMDARs) comprise a family of ligand-gated ionotropic glutamate receptors that mediate a slow, calcium-permeable component to excitatory neurotransmission. The GluN2D subunit is enriched in GABAergic inhibitory interneurons in cortical tissue. Diminished levels of GABAergic inhibition contribute to multiple neuropsychiatric conditions, suggesting that enhancing inhibition may have therapeutic utility, thus making GluN2D modulation an attractive drug target. Here, we describe the actions of a GluN2C/GluN2D-selective positive allosteric modulator (PAM), (+)-EU1180-453, which has improved drug-like properties such as increased aqueous solubility compared to the first-in-class GluN2C/GluN2D-selective prototypical PAM (+)-CIQ. (+)-EU1180-453 doubles the NMDAR response at lower concentrations (< 10 µM) compared to (+)-CIQ, and produces a greater degree of maximal potentiation at 30 µM. Using *in vitro* electrophysiological recordings, we show that (+)-EU1180-453 potentiates triheteromeric NMDARs containing at least one GluN2C or GluN2D subunit, and is active at both exon5-lacking and exon5-containing GluN1 splice variants. (+)-EU1180-453 increases glutamate efficacy for GluN2C/GluN2D-containing NMDARs by both prolonging the deactivation time and potentiating the peak response amplitude. We show that (+)-EU1180-453 selectively increases synaptic NMDAR-mediated charge transfer onto P11-15 CA1 *stratum radiatum* hippocampal interneurons, but is without effect on CA1 pyramidal cells. This increased charge transfer enhances inhibitory output from GABAergic interneurons onto CA1 pyramidal cells in a GluN2D-dependent manner. (+)-EU1180-453 also shifts excitatory-to-inhibitory coupling towards increased inhibition and produces enhanced gamma band power from carbachol-induced field potential oscillations in hippocampal slices. Thus, (+)-EU1180-453 can enhance overall circuit inhibition, which could prove therapeutically useful for the treatment of anxiety, depression, schizophrenia, and other neuropsychiatric disorders.

**Significance Statement:** Interneuron dysfunction and diminished GABAergic inhibition in neocortical and hippocampal circuits remains a prominent molecular hypothesis for neuropsychiatric diseases including anxiety, depression, and schizophrenia. Pharmacological agents that boost GABA receptor function have shown utility in various forms of depression and treating symptoms of schizophrenia. Cortical GABAergic interneurons, unlike their excitatory pyramidal cell counterparts, are enriched for the GluN2D subunit of the NMDA receptor. Thus, GluN2D subunit-selective modulation could be a useful therapeutic tool to enhance local inhibition, improving the prognosis for neuropsychiatric diseases for which interneuron dysfunction is prominent and causal to circuit aberration.

## Introduction

N-methyl-D-aspartate receptors (NMDARs) comprise a family of ligand-gated ionotropic glutamate receptors that mediate a calcium-permeable component to excitatory neurotransmission (Hansen et al., 2021). Glutamate-binding NMDARs are heterotetrameric assemblies, traditionally composed of two obligate GluN1 subunits (encoded by the *GRIN1* gene) and any combination of two GluN2A-D subunits encoded by the *GRIN2A*-*GRIN2D* genes (Hansen et al., 2021). Given their ubiquitous expression, participation in glutamatergic neurotransmission, and facilitation of calcium entry into cells, NMDARs have been implicated in a host of physiological and developmental roles including learning, memory, spatial navigation, coordinated movement, decision-making, neuronal migration, morphological development, and synaptic connectivity (Adesnik et al., 2008; Collingridge, 1987; Hansen et al., 2021; Komuro and Rakic, 1993; Nakazawa et al., 2004; Nash and Brotchie, 2000; Ultanir et al., 2007; Wang, 2002; Watanabe et al., 1998).

Modulation of NMDARs for therapeutic gain was initially pursued following the emergence of the NMDAR-mediated excitotoxicity hypothesis for neurodegenerative diseases including Alzheimer’s, Huntington’s, and Parkinson’s disease as well as acute injury associated with ischemic stroke, subarachnoid hemorrhage, hypoxia, or traumatic brain injury (Haichen Wang, 2024; Hoyte et al., 2004; Kemp and McKernan, 2002; Lipton, 2004; Lipton and Nicotera, 1998; Lipton and Rosenberg, 1994; Myers et al., 2021; Wang et al., 2014; Yuan et al., 2015). Although the first generation of non-selective NMDAR inhibitors were protective against excitotoxicity in multiple preclinical models, they also inhibited the channel’s ability to operate normally, which led to undesirable on target side effects such as drowsiness, hallucinations, motor dysfunction, sedation, and even coma, all of which complicate clinical care in emergency centers (Hoyte et al., 2004; Lipton, 2004). The failure of these compounds in the clinic had multiple underlying causes, including off-target liabilities, poorly defined endpoints, administration after the effective time window for neuroprotection, patient heterogeneity plus unfavorable on-target effects that led to dose-lowering (Myers et al., 2021). Early NMDAR inhibitors were either competitive antagonists (*e.g.*, Selfotel and analogs) or pore blockers such as Aptiganel (Cerestat), MK-801, memantine, dextromethorphan, and its metabolite dextrorphan. These inhibitors can produce complete block of all NMDARs, compromising the endogenous signaling ability of NMDARs within the circuit and blunting of basal NMDAR activity required for normal brain function. Moreover, these classes of inhibitors lacked strong NMDAR subunit specificity in that they were efficacious on GluN2A, GluN2B, GluN2C, and GluN2D-containing NMDARs. Thus, all NMDARs within the CNS were inhibited, including those with regulatory functions necessary for cardiovascular and motor control (Lipton, 2004).

Allosteric modulators of NMDARs can diminish some of the problems associated with traditional pore blockers and competitive antagonists by fine-tuning channel activity instead of producing complete block. Since allosteric modulators bind to less conserved regions of the receptor, rather than the highly conserved agonist binding pocket or channel pore (Hansen et al., 2021), allosteric modulators exhibit strong subunit selectivity. Ifenprodil and its derivatives were the first subunit-selective negative allosteric modulators (NAMs) developed for NMDARs, and showed promising pre-clinical efficacy in numerous indications, including pain (Chizh et al., 2001) and acute injury (Yuan et al., 2015). Studies in humans showed potential utility for treatment-resistant depression (Preskorn et al., 2008). These early-stage GluN2B selective NAMs such as eliprodil failed clinical trials, partly due to poor study design, as well as undesirable off- and on-target side-effects (Gogas, 2006; Herring et al., 2017; Ikonomidou and Turski, 2002; Mony et al., 2009).

Allosteric modulation of GluN2D-containing NMDARs has recently become appreciated as an attractive target due to their unique cell-type specific expression profile in the CNS. Although highly expressed during embryogenesis and early neonatal development, GluN2D expression in the developing and mature brain shows expression in cortical GABAergic interneurons (Perszyk et al., 2016; von Engelhardt et al., 2015), as well as select neurons in the spinal cord (Tolle et al., 1993), thalamus (O’Hara et al., 1995), olfactory bulb (Akazawa et al., 1994), basal ganglia (Swanger et al., 2015), and cerebellum (Akazawa et al., 1994). GluN2D hypofunction hypotheses have been suggested for both schizophrenia (Lisman et al., 2008; Schmitt et al., 2010) and Parkinson’s disease (Tozzi et al., 2016), with pre-clinical models suggesting that GluN2D PAMs can modulate prepulse inhibition and dopamine release from basal ganglia neurons (Nouhi et al., 2018; Suryavanshi et al., 2014), respectively. Disinhibition of medial ganglionic eminence (MGE)-derived interneurons such as somatostatin-positive (Fuchs et al., 2017; Salimando et al., 2020) and parvalbumin-positive (Volitaki et al., 2024) interneurons produce anxiolytic and antidepressant-like phenotypes in rodent models, suggesting that alteration of interneuron output is sufficient to modulate neuropsychiatric states. Within the bed nucleus of the stria terminalis (BNST), GluN2D-containing NMDAR signaling is implicated in anxiety- and depressive-like states (Salimando et al., 2020). Utilizing whole exome sequencing, *GRIN2D* human patient disease-associated *de novo* mutations have been attributed to the development of epilepsy, schizophrenia, bipolar disorder, autism, and intellectual disability (Camp and Yuan, 2020). Taken together, aberrant GluN2D-mediated signaling, particularly when viewed in the context of GABAergic interneuron function, is involved in the modulation of several neuropsychiatric disorders, highlighting the potential for GluN2D-targeting therapeutics.

(+)-(3-chlorophenyl)(6,7-dimethoxy-1-((4-methoxyphenoxy)-methyl)-3,4-dihydroisoquinolin-2(1*H*)-yl)methanone), referred to as (+)CIQ, was the first GluN2C/GluN2D-selective PAM described that showed strong selectivity over GluN2A- and GluN2B-containing NMDARs (Mullasseril et al., 2010). Identification of this class of modulator has enabled significant progress towards understanding the effects of GluN2C/GluN2D modulation in circuits and *in vivo* (Nouhi et al., 2018; Suryavanshi et al., 2014). However, CIQ’s potency and solubility are low, limiting utility as a tool compound. Recently, a new derivative, (+)-EU1180-453, has been synthesized with a 15-fold improvement in doubling concentration (*i.e.*, the concentration needed to double a current response) and more than a 7-fold improvement in aqueous solubility (Epplin et al., 2020). We examine here the effects of (+)-EU1180-453 on the NMDAR response time course, efficacy on triheteromeric NMDARs that contain two different GluN2 subunits, and on synaptic NMDARs. We also assessed the ability of (+)-EU1180-453 to enhance GABAergic interneuron network function, which could provide data to support future pre-clinical studies that investigate GluN2D-selective PAMs as therapeutic agents.

## Methods

### Novel Compounds

The synthesis of (+)-EU1180-453 has been previously described (Epplin et al., 2020). Stock solutions of (+)-EU1180-453 were made fresh on a weekly basis and dissolved in DMSO. Aliquots were made in concentrations of 0 µM (vehicle) or 10 mM and stored at -20C. Stock solutions were thawed only once for use and discarded; no aliquots were ever refrozen.

### Molecular Biology

All cDNAs encoding NMDAR subunits were from rat. cDNAs encoding GluN1 lacking exon5 (GluN1-1a; GenBank U11418), GluN1 containing exon5 (GluN1-1b; GenBank U08261), GluN2A (Gen Bank <u>D13211</u>), GluN2B (GenBank <u>U11419</u>), GluN2C (GenBank <u>M91563</u>), and GluN2D (GenBank <u>L31611</u>) were provided by Drs. S. Heinemann (Salk Institute, La Jolla, CA), S. Nakanishi (Osaka Bioscience Institute, Osaka, Japan), and P. Seeburg (Max Planck Institute for Medical Research, Heidelberg, Germany). Site-directed mutagenesis was performed using the QuikChange strategy (Agilent Technologies). Amino acids were numbered according to the full-length protein, with the inclusion of the signal peptide (the initiating methionine is 1). For the synthesis of cRNA, cDNA constructs were first linearized using the appropriate restriction enzymes, purified using a QIAquick purification kit (Qiagen, Germantown MD), and then utilized for *in vitro* transcription according to the manufacturer’s instructions (mMessage mMachine, Ambion, Austin, TX).

Chimeric cDNA constructs that enable user control of subunit stoichiometry were synthesized using GluN2A C-terminal with in frame peptide tags encoding coiled-coil domains and an endoplasmic reticulum (ER) retention signal, as previously described (Hansen et al., 2014). Two peptides comprising a synthetic helix, leucine zipper motifs from GABA_B1_ (referred to as C1) or GABA_B2_ (referred to as C2), and a di-lysine KKTN ER retention signal were inserted in-frame in place of the stop codon for GluN2A to yield GluN2A_C1_ and GluN2A_C2_ (Jackson et al., 1990, 1993; Zerangue et al., 2001). This only permits NMDARs with one copy of a subunit with a C1 tag and one copy of a subunit with a C2 tag to mask the ER retention signal and reach the cell surface, allowing us to express triheteromeric NMDARs that contain two different GluN2 subunits (Hansen et al., 2014). The C-terminal tail of the GluN2B, GluN2C, and GluN2D subunits (residues after 838 for GluN2B, 837 for GluN2C, and 862 for GluN2D) were replaced by the modified C-terminal domain of GluN2A_C1_ or GluN2A_C2_ (residues after 837 in GluN2A). To experimentally determine whether any diheteromeric NMDARs reached the cell surface, we generated constructs with two mutations in the glutamate binding site (referred to as RK/TI) of each GluN2 subunit (GluN2A-R518K,T690I; GluN2B-R519K,T691I; GluN2C-R529K,T701I; GluN2D-R543K,T715I). This allowed determination of the ability of the ER retention signal to inhibit trafficking to the cell surface. C1–C1-expressing receptors (diheteromeric receptor) will generate a current response that can be isolated and recorded when the C2-containing subunit has the RK/TI mutations (Yi et al., 2017), and similarly, C2-C2-expressing receptors can be recorded in isolation when the C1-containing subunit has the RK/TI mutations. All constructs were verified by sequencing. Wild-type and modified GluN2 cDNAs were subcloned into pCI-neo vectors (Promega, Madison, WI) and used for *in vitro* cRNA for expression in *Xenopus* oocytes. GluN1 was subcloned into a pGEM-HE vector.

### Two Electrode Voltage Clamp Recordings from Xenopus Oocytes

Defolliculated *Xenopus laevis* oocytes (stage V–VI) were purchased from Ecocyte BioScience (Austin, TX), or obtained from Xenopus-1 and isolated as previously described (Xie et al. 2023). Oocytes were injected with cRNAs for wild-type GluN1 and GluN2 at a 1:2 ratio in a final volume of 50 nL (0.2–10 ng of total cRNA was used). cRNAs were diluted with RNase-free water and generated responses ranging from 200 to 2000 nA. After cRNA injection, the oocytes were incubated at 15 °C in Barth’s solution containing (in mM) 88 NaCl, 2.4 NaHCO_3_, 1 KCl, 0.33 Ca(NO_3_)_2_, 0.41 CaCl_2_, 0.82 MgSO_4_, 5 HEPES, pH 7.4 with NaOH, supplemented with 100 U/mL penicillin, 100 *μ*g/mL streptomycin (Invitrogen, Carlsbad, CA), and 100 *μ*g/mL gentamicin (Fisher Scientific, Pittsburgh, PA). For recordings with triheteromeric receptors, oocytes were incubated at 19 °C, except GluN1/GluN2B_C1_/GluN2D_C2_ and GluN1/GluN2A_C1_/GluN2D_C2_, which were incubated at 15 °C for 1–2 h and then at 19 °C until recording.

Recordings on cRNA-injected oocytes were performed 2–5 days following injection at room temperature (23 °C) using a two-electrode voltage-clamp amplifier (OC725, Warner Instrument, Hamilton, CT). Responses were elicited by bath-application of saturating concentrations glutamate (100 *μ*M) and glycine (30 *μ*M), unless otherwise stated. Responses were low-pass filtered at 10–20 Hz (4-pole, −3 dB Bessel; Frequency Devices, Haverhill, MD) and digitized at the Nyquist rate using PCI-6025E or USB-6212 BNC data acquisition boards (National Instruments, Austin, TX). Oocytes were housed in a custom-made recording chamber and continuously perfused (∼2.5 mL/min) with recording solution containing (in mM) 90 NaCl, 1 KCl, 10 HEPES, 0.5 BaCl_2_, 0.01 EDTA (pH 7.4 with NaOH). Solutions were applied by gravity, and solution exchange was controlled through a digital 8-valve modulator positioner (Hamilton, Reno, NV). Recording electrodes contained 0.3–3.0 M KCl, and responses were recorded at a V_hold_ −40 mV. Data acquisition, voltage control, and solution exchange were controlled by custom software.

Triheteromeric NMDAR expression was achieved by injecting into oocytes varying amounts of cRNA in 50 nL of water. Injections for GluN1/GluN2A_C1_/GluN2C_C2_ were performed as previously described in (Bhattacharya et al., 2018). Injections for GluN1/GluN2A_C1_/GluN2B_C2_ were performed as previously described in (Hansen et al., 2014); 50 nl of 40 mM BAPTA was injected at time of recording to limit gradual increases in the response due to divalent activated second messenger mechanisms, which we refer to as run-up. Injections for GluN1/GluN2B_C1_/GluN2D_C2_ were performed as previously described in (Yi et al., 2019); 50 nl of 40 mM BAPTA was injected at time of recording to limit response run-up. GluN1/GluN2A_C1_/GluN2D_C2_ cRNAs were co-injected at ratio of 1:1:4 (total cRNA was 1.5 ng in 50 nL of RNase free water) 50 nl of 40 mM BAPTA was injected at time of recording to limit response run-up. For all triheteromeric experiments, the cRNA ratios for injection for the RK+TI mutations and diheteromeric controls matched that of the triheteromeric construct. On average, co-expression of GluN1/GluN2A_C1_/GluN2B_C2_ showed a summed escape current less than 7%, co-expression of GluN1/GluN2A_C1_/GluN2C_C2_ and GluN1/GluN2B_C1_/GluN2D_C2_ showed a summed escape current of less than 10%, and co-expression of GluN1/GluN2A_C1_/GluN2D_C2_ showed a summed escape current of less than 9%.

### Whole-Cell Patch-Clamp Recordings from HEK293T Cells

HEK293T cells (CRL 1573, ATCC, Rockville, MD, hereafter HEK cells) were cultured in Dulbecco’s modified Eagle medium with GlutaMax-I (Invitrogen, Carlsbad, CA), 10% dialyzed fetal bovine serum, 10 U/mL penicillin, and 10 *μ*g/mL streptomycin at 37 °C in an atmosphere of 5% CO_2_. HEK cells were plated in 24-well plates on 5 mm glass coverslips (Warner Instruments, Hamden, CT) coated with 10–100 *μ*g/mL poly(d-lysine). Cells were transfected with GluN1, GluN2, and GFP in plasmids at a ratio of 1:1:1 using the calcium phosphate method (Chen and Okayama, 1987). 10 *μ*L of plasmid DNA in water (0.2 *μ*g/*μ*L; total 0.5 *μ*g of DNA per well) was mixed with 65 *μ*L of H_2_O and 25 *μ*L of 1 M CaCl_2_ and left to incubate for 3-5 minutes. Then, this mixture was added to 100 *μ*L of 2× BES solution composed of (in mM) 50 *N*,*N*-bis(2-hydroxyethyl)-2-aminoethanesulfonic acid (BES), 280 NaCl, and 1.5 Na_2_HPO_4_ (adjusted to pH 6.95 with NaOH). After another 5–10 minutes, 50 *μ*L of the resulting solution was added dropwise to each of four wells of a 24-well plate containing HEK cells and 0.5 mL of DMEM. The culture medium containing the transfection mixture was replaced after 4-6 hours with DMEM supplemented with penicillin, streptomycin, and the NMDAR antagonists D,L-2-amino-5-phosphonovalerate (200 *μ*M) and 7-chlorokynurenate (200 *μ*M). Experiments were performed 12–24 h post-transfection.

Whole-cell voltage-clamp recordings were performed at V_hold_ −60 mV at room temperature (23 °C). Current responses were recorded using an Axopatch 200B amplifier (Molecular Devices, Union City, CA), low-pass filtered at 4–8 kHz (−3 dB, 8-pole Bessel; Frequency Devices, Haverhill, MD), and digitized at 40 kHz using Digidata 1322A or 1440A (Molecular Devices, Union City, CA) controlled by Clampex software (Molecular Devices, Union City, CA). Recording electrodes (3–4 MΩ) were pulled from thin-walled borosillicate glass (TW150F-4; World Precision Instruments, Sarasota, FL) and filled with (in mM) 110 D-gluconic acid, 110 CsOH, 30 CsCl, 5 HEPES, 4 NaCl, 0.5 CaCl_2_, 2 MgCl_2_, 5 BAPTA, 2 NaATP, and 0.3 NaGTP (pH 7.35 with CsOH). The extracellular solution contained (in mM) 150 NaCl, 10 HEPES, 3 KCl, 0.5 CaCl_2_, 0.01 EDTA, and 20 D-mannitol (adjusted to pH 7.4 with NaOH). Rapid solution exchange was accomplished through a two-barrel theta-glass controlled by a piezoelectric translator (Burleigh Instruments, Fishers, NY). Open tip junction currents using undiluted and diluted (1:2 in water) extracellular recording solution were used to estimate the speed of solution exchange at the tip, which had 10–90% rise times of 0.4–0.8 ms.

Transfected HEK cells were recorded under whole-cell voltage-clamp (V_hold_ -70 mV), physically lifted from glass coverslips, and placed in front of a two-barrel theta tube; the position (and flow pipe the cell was exposed to) was driven by a piezoelectric translator. To start, all cells were continuously bathed in saturating glycine (100 µM) and then rapidly exposed to saturating concentrations of both glutamate (1 mM) and glycine (100 µM) for 10 ms, then moved back into saturating glycine (100 µM). This time course of co-agonist exposure is designed to mimic synaptically-released glutamate and thus provide an estimate of a synaptic-like current. After baseline responses were established, cells were then exposed to saturating glycine (100 µM) plus 10 µM (+)-EU1180-453, rapidly exposed to saturating concentrations of both glutamate (1 mM) and glycine (100 µM) plus 10 µM (+)-EU1180-453 for 10 ms, then moved back into saturating glycine (100 µM) plus 10 µM (+)-EU1180-453. Experiments were finished by removing (+)-EU1180-453 from all tubes and re-establishing a baseline current with jumps into saturating glutamate (1 mM) for 10 ms in the constant presence of saturating glycine (100 µM).

Experiments started by recording 15-20 responses to 1.5 second glutamate application (20-60 second intervals) followed by 15-20 responses to brief 10 ms application of glutamate (10-30 second intervals). Then, 10 µM (+)-EU1180-453 was washed onto the cell for 3 minutes by adding it to both sides of the theta tube. After 3 minutes, 15-20 responses to 1.5 second application of glutamate (20-60 second intervals) plus 10 µM (+)-EU1180-453 were recorded followed by 15-20 responses to 10 ms application of glutamate (10-30 second intervals) plus 10 µM (+)-EU1180-453. 10 µM (+)-EU1180-453 was then washed out of the theta tube for 3 minutes and ‘recovery’ measurements were obtained by recording 15-20 responses to 1.5 second application of glutamate (20-60 second intervals) followed by 15-20 responses to 10 ms application of glutamate (10-30 second intervals).

### Animals and Breeding

All mouse procedures were conducted at Emory University, which is fully accredited by AAALAC, approved by the Emory IACUC, and performed in accordance with state and federal Animal Welfare Acts and Public Health Service policies. C57BL/6J mice from the Jackson Laboratory (catalog number: 000664) were used as wild type (WT) mice. *Grin2d*^-/-^ mice (homozygous, global *Grin2d*-knockout mice (Ikeda et al., 1995) were obtained from RIKEN, through the National Bio-Resource Project of the MEXT, Japan and were backcrossed > 10 times onto a C57BL/6J background. Juvenile postnatal day (P) 11-15 of both sexes were used in all experiments. Mice were housed in ventilated cages at controlled temperature (22–23°C), humidity ∼60%, and had *ad libitum* access to regular chow and water. WT mice were on a 12 h light:dark cycle; *Grin2d*^-/-^ mice were on a 10:14 h light:dark cycle. *Grin2d*^-/-^ mice were also kept away from heavy foot traffic and experienced minimal human interaction to help promote increased breeding habits.

### Acute Hippocampal Slice Preparation and Electrophysiological Recordings

Mouse hippocampal slices were prepared as previously described (Camp et al., 2023). Mice were overdosed with inhaled isoflurane, brains were rapidly removed and immediately placed in an ice-cold, sucrose-based artificial cerebrospinal fluid (aCSF) containing the following (in mM): 88 sucrose, 80 NaCl, 2.5 KCl, 1.25 Na_2_HPO4, 26 NaHCO_3_, 10 glucose, 2 thiourea, 3 sodium pyruvate, 5 sodium ascorbate, 12 N-acetylcysteine, 10 MgSO_4_, and 0.5 CaCl_2_ bubbled in 95% O_2_/5% CO_2_. 300-µm thick horizontal hippocampal sections were obtained using a vibratome (Lecia, VT1200S). After sectioning, slices were incubated in the sucrose-based aCSF as described above but with 4 mM MgSO_4_ at 32°C for 30 minutes then returned to room temperature for at least an hour before use. All cells were visualized using an upright Olympus BX50W microscope with IR-DIC optics coupled to a Dage IR-2000 camera. Whole-cell patch clamp recordings were obtained using a Multiclamp 700B (Molecular Devices, digitized at 20 kHz using a Digidata 1440a (Molecular Devices) controlled by pClamp 10.6 software (Molecular Devices). All signals were low-pass filtered at 2 kHz using a Bessel 8-pole filter (-3 dB, Warner, LPF-8).

For all voltage-clamp experiments, the following intracellular solution was used (in mM): 100 Cs-gluconate, 5 CsCl, 0.6 EGTA, 5 BAPTA, 5 MgCl_2_, 8 NaCl, 2 Na-ATP, 0.3 Na-GTP, 40 HEPES, 5 Na-phosphocreatine, and 3 QX-314 supplemented with 0.5% biocytin. Evoked NMDAR-mediated EPSC recordings were made in the following aCSF extracellular solution (in mM): 126 NaCl, 2.5 KCl, 1.25 Na_2_HPO_4_, 26 NaHCO_3_, 20 glucose, 0.2 MgSO_4_, and 1.5 CaCl_2_ bubbled with 95% O_2_/5% CO_2_. sIPSC and EPSP/IPSP data were obtained with the same extracellular solution, except with 1.5 mM Mg^2+^. All voltage-clamp experiments were performed at 30-32°C using an inline heater (Warner, SH-27B).

For evoked NMDAR-mediated EPSCs, a monopolar iridium-platinum stimulating electrode (FHC, Inc.) was placed in the upper 1/3^rd^ of the Schaffer collaterals within CA1 *stratum radiatum* to elicit a single 50 µs stimulation at a frequency of 0.03 Hz. The NMDAR-mediated EPSC was pharmacologically isolated with 10 μM NBQX and 10 μM gabazine. Cells were held at -30 mV and stimulation intensity was chosen to be near 50% of the maximum peak amplitude of the NMDAR-mediated EPSC. Baseline recordings were made for a total of five minutes. Then DMSO or 10 µM (+)-EU1180-453 was washed onto the slice for ten minutes and response recordings were made for a total of five minutes. A total of 8-12 epochs were obtained and averaged together for each baseline and response period. At the conclusion of the recording, 200 μM DL-APV for pyramidal cells or 400 μM DL-APV for *stratum radiatum* interneurons was applied to ensure that the responses were mediated via NMDARs.

For sIPSCs, cells were held at +10 mV sIPSCs, which is the reversal potential for ionotropic glutamate receptors. Baseline recordings were made for five minutes, and then DMSO or 10 µM (+)-EU1180-453 was washed onto the slice for ten minutes and response recordings were made for five minutes. All experiments ended by washing on 10 µM gabazine to ensure that the recorded responses were mediated by GABA_A_ receptors.

For EPSP/IPSP recordings, CA1 pyramidal cells were current-clamped and held at -60 mV using the following internal solution, in mM: 115 potassium gluconate, 0.6 EGTA, 2 MgCl_2_, 2 Na_2_ATP, 0.3 Na_2_GTP, 10 HEPES, 5 sodium phosphocreatine, 8 KCl, and 0.3-0.5% biocytin (pH adjusted to 7.35 using CsOH). If injected current required to maintain juvenile CA1 pyramidal cells at -60 mV exceeded ± 50 pA, the cell was rejected. Schaffer collateral stimulation (0.03 Hz, 50 µs stimulation duration) was set 10-20% below threshold to generate a single action potential, usually 50-100 µA. Baseline recordings were obtained for 5 minutes and 10 µM (+)-EU1180-453 or 0.1% DMSO was applied for 10 minutes.

For carbachol-induced gamma oscillations, 400-um thick slices were used. Field potential recordings from *stratum pyramidale* in CA1 were recorded using an interface-style chamber. External solution was composed of the following (in mM): 126 NaCl, 3 KCl, 1.25 Na_2_HPO_4_, 26 NaHCO_3_, 20 glucose, 1.5 MgSO_4_, and 2.5 CaCl_2_ bubbled with 95% O_2_/5% CO_2_. External solution was heated to 30-32°C using an inline heater (Warner, SH-27B). Field potential electrodes were filled with external solution. Data were obtained using an A-M Systems Model 1800 field recording amplifier using 1,000× gain, with low and high pass filtering at 8 and 200 Hz, respectively. Recordings were digitized using a National Instruments PCI-6210 board and recorded at 10 kHz using NACGather software (Theta Burst Corporation, 2001). Slices were placed into the chamber and allowed to equilibrate for 15-20 minutes. Baseline periods were recorded for five minutes, 20 µM carbachol was applied for 15 minutes, and then 20 µM carbachol plus 10 µM (+)-EU1180-453 was applied for another 10 minutes. The last minute of each period was analyzed for power. If total baseline power (0-100 Hz) exceeded 2.0e^-4^ μV2/Hz, power in 20 µM carbachol only increased power at 60 Hz (noise), or total gamma power in 20 µM carbachol did not exceed 4-fold over baseline gamma power, the slice was excluded.

For all electrophysiological recordings, a wash-in period of 10-15 minutes was used with (+)-EU1180-453 application as previous experiments showed this time was sufficient to achieve maximal and steady-state levels of drug action. After each experiment, the perfusion lines were washed with 70% ethanol and then deionized H_2_0 to remove any remaining compound.

### Data Analysis, Statistical Tests, and Figure Preparation

For oocyte experiments, concentration-response data was analyzed in GraphPad 5.0. Responses evoked by test compounds are expressed as a percentage of the initial response to glutamate and glycine alone. Data for individual cells were fit with the Hill equation:

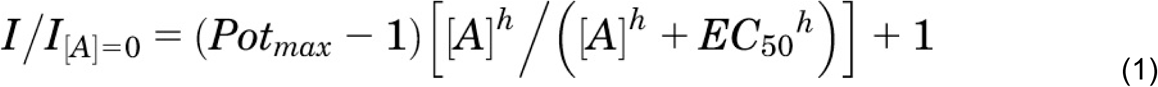

where *I* is the current response, *Pot_max_* is the maximal predicted potentiation of the glutamate/glycine response (and expressed in the manuscript as a percent for clarity), [*A*] is the concentration of the modulator, *h* is the Hill slope, and *EC_50_* is the half-maximally effective concentration of the modulator. The potentiated current response was expressed as percent of control. Displayed fitted curves were obtained by simultaneously fitting all oocyte data from each concentration.

For all whole-cell current responses, all data were analyzed offline using ChanneLab (SynaptoSoft) or pClamp 10.3 (Molecular Devices) on the average response per condition (i.e. baseline period, response period, recovery period) except for charge transfer for evoked NMDAR-mediated EPSCs, where each trace was analyzed individually. Rise time was measured as the 10-90% fractional response of the rising phase of each current response. Peak amplitude was determined by finding the maximum of a moving search average of 8-12 data points. Desensitization in HEK cells was measured as the fraction of steady-state amplitude (I_SS_) ∼100 ms before the end of application period and dividing by the peak amplitude (I_peak_, amplitude at the beginning of application period). Deactivation for HEK cells and evoked NMDAR-mediated EPSCs were measured using a dual-exponential function and reported as weighted deactivation time constants (τ_W_), calculated using the following equation:

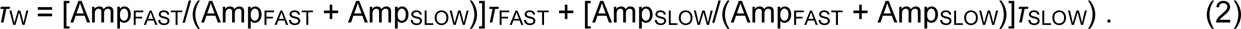

Charge transfer of current responses was calculated as the area under the curve (amplitude*tau) for each response.

sIPSC data were analyzed offline using MiniAnalysis (Synaptosoft) with an 8-pA amplitude threshold for event detection, which was 5-fold higher than our peak RMS noise. Decay time for sIPSC responses were the average of 20-30 randomly chosen responses throughout each condition (i.e. baseline vs drug application). A single exponential function was fitted to each composite trace. EPSP/IPSP peak amplitudes were analyzed offline using Clampfit 10.3 (Molecular Devices). Any traces with action potential generation or lack of clear separation from the stimulus artifact were excluded. Carbachol-induced oscillations were analyzed offline using Clampfit 10.3 (Molecular Devices). One-minute stretches of data were first post-hoc low- and high-pass filtered at 5 Hz (Bessel, 8-pole) and 120 Hz (Gaussian). Then, using the power spectrum feature in Clampfit, the Blackman algorithm was selected and used at 8192 bits (2.4 Hz of spectral resolution). Power bands were selected as follows: 0-4 Hz (delta), 4-8 Hz (theta), 8-12 Hz (alpha), 12-20 Hz (beta), and 20-100 Hz (gamma). Since recordings were made below physiological temperatures of 37°C, the gamma band window was shifted from 30-100 Hz to 20-100 Hz as previously described (Dickinson et al., 2003; Mackenzie-Gray Scott et al., 2022).

Series resistance was monitored throughout all experiments and was typically 8−22 MΩ. For voltage clamp recordings, a 50 ms duration 5-mV square wave was included in the stimulation paradigm (once every 30 seconds) or inserted once every 30 seconds in gap-free mode for sIPSCs. Series resistance was monitored throughout the recording and determined offline by analyzing the peak of the capacitive charging spike and applying Ohm’s law. If the series resistance changed >25% during the experiment, or ever exceeded 30 MΩ, the cell was excluded.

Paired Student’s t-test (two-tailed), one-way ANOVA, two-way ANOVA, and the Kolomorgov-Smirnov tests were used when appropriate. For experiments where multiple statistical analyses were performed on the same dataset, our significance threshold was lowered to correct for family-wise error rate (FWER) using the Bonferroni post-hoc correction method. All studies were designed so that an effect size of at least 1 was detected at 80% or greater power. All statistical analyses were performed in Prism’s GraphPad software all figures were generated in Adobe’s Illustrator software.

## Results

### (+)-EU1180-453 Potentiates Triheteromeric GluN2C- and GluN2D-Containing NMDARs

(+)-CIQ was the first-in-class GluN2C/GluN2D-selective PAM of NMDARs. Its chemical structure contained a tetrahydroisoquinoline scaffold, which was replaced by a 6-5 heterocyclic ring system to generate (+)-EU1180-453 (**Figure 1A**) (Epplin et al., 2020). These modifications resulted in an increase in both potency (EC_50_ 8.9 µM for (+)-CIQ vs EC_50_ 3.2 µM for (+)-EU1180-453 at GluN2D) and maximal achievable potentiation at saturating concentrations (350% over baseline for both GluN2C and GluN2C) (Epplin et al., 2020). Unlike (+)-CIQ (Mullasseril et al., 2010), saturating concentration of (+)-EU1180-453 (30 µM) increases glutamate potency at both GluN2C-(0.80 µM without (+)-EU1180-453 vs 0.31 µM with (+)-EU1180-453; **Figure 1B**) and GluN2D-containing NMDARs (0.53 µM without (+)-EU1180-453 vs 0.25 µM with (+)-EU1180-453; **Figure 1C**). Together these enhancements reduced the doubling concentration (31.6 µM for (+)-CIQ vs 2.0 µM for (+)-EU1180-453 for GluN1/GluN2D), which is the concentration required to increase the basal response without the modulator to saturating concentrations of agonist (100 µM glutamate/30 µM glycine) by 2-fold (Epplin et al., 2020). The aqueous solubility was also increased over 9-fold, from 8 µM for (+)-CIQ to 74 µM for (+)-EU1180-453, while still possessing favorable brain penetration and half-life suitable for acute *in vivo* preclinical experiments (Epplin et al., 2020).

**Figure 1.**
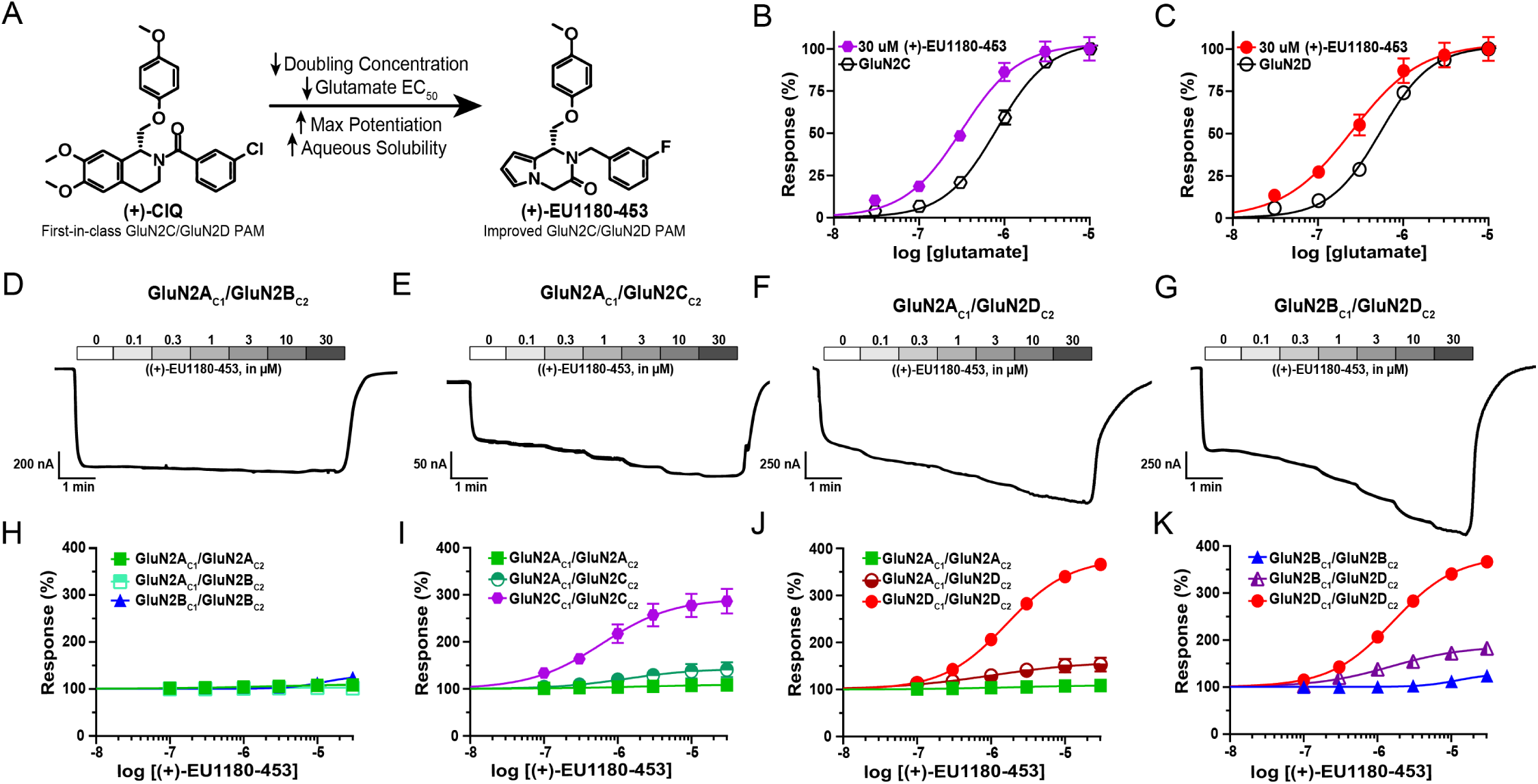
(+)-EU1180-453 potentiates response of GluN2C- and GluN2D-containing triheteromeric NMDARs expressed in *Xenopus* oocytes. **A)** Chemical structure of (+)-CIQ and (+)-EU1180-453. Response of **B)** GluN2C-containing NMDARs and **C)** GluN2D-containing NMDARs to varying concentrations of glutamate in presence of 30 µM glycine and 30 µM (+)-EU1180-453. Representative current responses of **D)** GluN2A_C1_/GluN2B_C2_, **E)** GluN2A_C1_/GluN2C_C2_, **F)** GluN2A_C1_/GluN2D_C2_, and **G)** GluN2B_C1_/GluN2D_C2_ triheteromeric NMDARs to varying concentrations of (+)-EU1180-453 in presence of saturating concentrations of glutamate (100 µM) and glycine (30 µM). Fitted concentration-response curves of **H)** GluN2A_C1_/GluN2B_C2_, **I)** GluN2A_C1_/GluN2C_C2_, **J)** GluN2A_C1_/GluN2D_C2_, and **K)** GluN2B_C1_/GluN2D_C2_ triheteromeric NMDARs to varying concentrations of (+)-EU1180-453 in presence of 30 µM glycine and 100 µM glutamate. Diheteromeric NMDARs with corresponding C1- and C2-tails were included as reference. Maximum potentiation of percent response (amplitude with saturating glycine and glutamate only) was **H)** 103 ± 2.2% of control for GluN2A_C1_/GluN2B_C2_, **I)** 141 ± 13% of control for GluN2A_C1_/GluN2C_C2_, **J)** 153 ± 16% of control for GluN2A_C1_/GluN2D_C2_, and **K)** 183 ± 5.6% of control for GluN2B_C1_/GluN2D_C2_. All data are mean ± SEM. Number of oocytes for each receptor type ranged from N=10-18, from 2-3 frogs.

Many NMDARs in the brain are thought to be triheteromeric receptors that contain two GluN1 subunits and two different GluN2 subunits (Bhattacharya et al., 2018; Hansen et al., 2014; Luo et al., 1997; Rauner and Kohr, 2011; Swanger et al., 2015; Tovar et al., 2013). To assess the ability of (+)-EU1180-453 to potentiate triheteromeric NMDARs, we utilized chimeric NMDAR constructs in which the C-terminus contained a synthetic linker, a coiled-coil domain, and unique endoplasmic reticulum (ER) retention sequence (see Methods). Together, these modifications permit only receptors that contain the two complementarily-tagged GluN2 subunits to reach the cell surface (Hansen et al., 2014). We investigated eight different NMDARs that contained GluN1 coexpressed with GluN2A_C1_/GluN2A_C2_, GluN2BC1/GluN2BC2, GluN2CC1/GluN2CC2, GluN2DC1/GluN2DC2, GluN2AC1/GluN2BC2, GluN2AC1/GluN2CC2, GluN2AC1/GluN2DC2, and GluN2BC1/GluN2DC2. Concentration-response relationships were evaluated using maximal concentrations of co-agonists (100 µM glutamate and 30 µM glycine) and seven different concentrations of (+)-EU1180-453 (ranging from 0 µM to 30 µM).

We observed no potentiation at saturating concentrations (30 µM) of (+)-EU1180-453 for GluN2A_C1_/GluN2A_C2_ and GluN2A_C1_/GluN2B_C2_ (**Figure 1D, H**). GluN2B_C1_/GluN2B_C2_ receptors showed very slight potentiation (maximal potentiation was 124 ± 2% of control) at 30 µM (+)-EU1180-453 with an EC_50_ of 19 µM (**Figure 1E,I**). GluN2A_C1_/GluN2C_C2_ showed modest potentiation (maximum potentiation was 141 ± 13% of control) at 30 µM (+)-1180-453 with an EC_50_ of 1.6 µM (**Figure 1F,J**). Triheteromeric GluN2B_C1_/GluN2D_C2_ receptors showed a near doubling in maximum potentiation (183 ± 5.6% of control) with an EC_50_ of 1.7 µM (**Figure 1G,K**). GluN2C_C1_/GluN2C_C2_ and GluN2D_C1_/GluN2D_C2_ NMDARs showed the strongest potentiation, with GluN2C_C1_/GluN2C_C2_ receptors reaching 287% of maximum response with an EC_50_ 0.7 µM (**Figure 1J**) and GluN2D_C1_/GluN2D_C2_ receptors reaching 357% of maximum response with an EC_50_ of 1.7 µM (**Figure 1K**). Taken together, these data indicate that (+)-EU1180-453 shows strong preference for GluN2C- and GluN2D-containing diheteromeric NMDARs, with modest but significant potentiation of NMDARs containing only one copy of GluN2C or GluN2D. See **Supplemental Table S1** for a summary of all fitted results.

### (+)-EU1180-453 Enhances Charge Transfer of GluN2C- and GluN2D-Containing NMDARs

Next, we evaluated (+)-EU1180-453’s impact on NMDAR-mediated charge transfer in transiently transfected HEK cells. We started with an agonist exposure time of 10 ms (see *Methods*), which is designed to mimic synaptically-released glutamate and thus provide an estimate of a synaptic-like current. We observed a significant, but minor potentiation in fold charge transfer for GluN2A expressing NMDARs (1.9 ± 0.3 fold change over baseline; **Figure 2A,E**), but no impact on charge transfer for GluN2B receptors (1.4 ± 0.1 fold change over baseline; **Figure 2B,E**). Fold change of change transfer for GluN2C receptors and GluN2D receptors was significant and robust, with 6.5 ± 0.9 for GluN2C (**Figure 2C,E**) and 4.7 ± 0.9 for GluN2D (**Figure 2D,E**). We observed no change in peak amplitude following 10 µM (+)-EU1180-453 for GluN2A or GluN2B NMDARs. Application of 10 µM (+)-EU1180-453 produced a small, but significant, prolongation of the weighted tau used to describe receptor deactivation for GluN2A NMDARs and GluN2B NMDARs (**Supplemental Figure S1A,B**). (+)-EU1180-453 was much more active for GluN2C- and GluN2D-containing NMDARs, producing robust and significant potentiation of peak amplitude, prolongation of the weighted tau, and overall charge transfer at both subunits (**Supplemental Figure S1C,D)**. See **Supplemental Table S2** for full details We also evaluated (+)-EU1180-453 for agonist-dependent activity. Since our paradigm involved applying a subsaturating concentration of modulator (10 µM) before receptor activation, we can use the 10-90% rise time of the current response as a readout for any agonist-dependency (Perszyk et al., 2018). We found no significant difference in the 10-90% rise time between baseline and (+)-EU1180-453 application for any NMDAR subunit (GluN2A, GluN2B, GluN2C, and GluN2D; **Supplemental Table S2**). Thus, (+)-EU1180-453 does not exhibit detectable agonist-dependence.

**Figure 2.**
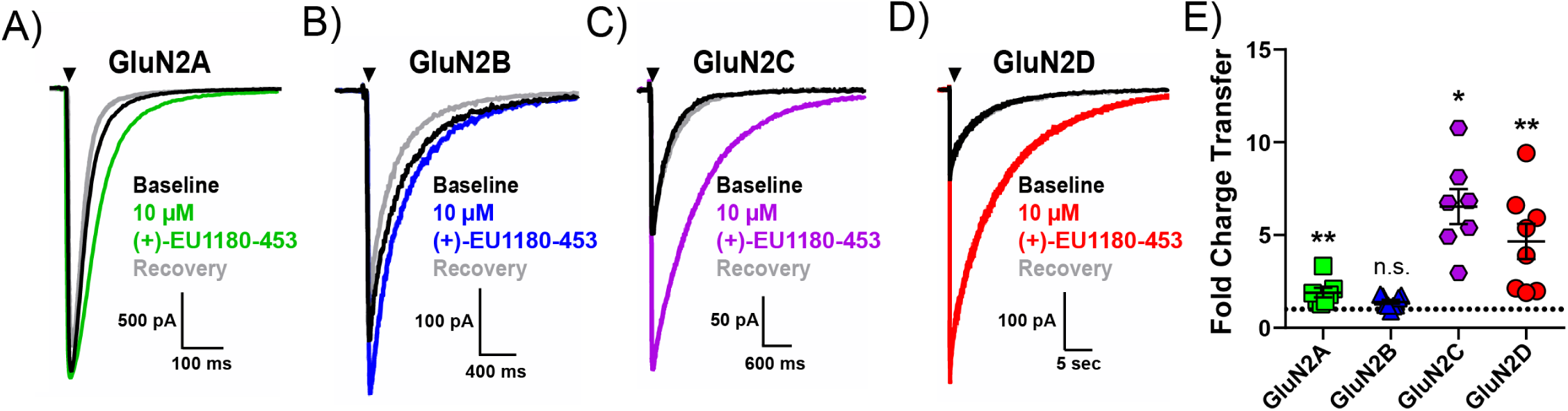
(+)-EU1180-453 strongly potentiates GluN2C- and GluN2D-containing NMDAR current responses recorded from HEK293T cells. Cells were held at -70 mV and subjected to rapid and brief (10 ms) application of saturating glutamate (1 mM) in the presence of 100 µM glycine. Current responses were recorded in absence, presence, and following removal of (+)-EU1180-453. Representative responses are shown for baseline (glutamate only, black trace), glutamate plus 10 µM (+)-EU1180-453 (color trace), and glutamate recovery (gray trace) for **A)** GluN1/GluN2A (n=7 cells), **B)** GluN1/GluN2B (n=9 cells), **C)** GluN1/GluN2C (n=7 cells), and **D)** GluN1/GluN2D (n=8 cells). **E)** The mean fold-change in charge transfer (response/baseline) is shown. The dotted line is at fold change of 1. All data are mean ± SEM. Statistical significance determined by paired t-test of control vs drug. See **Table 2** for full details. * p < 0.05; ** p < 0.01; n.s. indicates not significant.

Similar trends were observed when the length of the glutamate or glutamate plus (+)-EU1180-453 application was prolonged to 1.5 seconds (**Supplemental Figure S2, Supplemental Table S3**). Using this prolonged application of glutamate or glutamate plus (+)-EU1180-453, we can also estimate the extent of receptor desensitization by approximating the steady-state amplitude (I_SS_) ∼100 ms before the end of the agonist application period and dividing by the peak amplitude (I_peak_, amplitude at beginning of application period). There were no meaningful changes in the extent of receptor desensitization for GluN2A-, GluN2B-, GluN2C-, or GluN2D-containing NMDARs (**Supplemental Table S3**). Using this 1.5 second application paradigm, (+)-EU1180-453 did not impact overall charge transfer for GluN2A NMDARs (**Supplemental Figure S2A,E**) with a minor increase in charge transfer for GluN2B NMDARs (**Supplemental Figure S2B,E**). As observed previously, overall charge transfer at both GluN2C- and GluN2D-containing NMDARs (**Supplemental Figure S2C,D,E**) were markedly and significantly increased after (+)-EU1180-453 application. These data further support the strong preference of (+)-1180-453 to potentiate GluN2C and GluN2D NMDARs.

### (+)-EU1180-453 Potentiates Exon5-Containing GluN2C and GluN2D NMDARs

Within native synapses, NMDARs can also contain different splice variants of the GluN1 subunit. Although there are 8 total GluN1 splice variants (Hansen et al., 2021), heterologous electrophysiological assays indicate the presence or absence of exon5 within the ATD of GluN1 being the only alternatively spliced exon to influence the receptor response time course (Rumbaugh et al., 2000; Vance et al., 2012). While exon5-lacking GluN1 subunits (referred to as GluN1-1a) are the predominantly expressed splice variant in the CNS, exon5-containing GluN1 subunits (referred to as GluN1-1b) are expressed in native synapses, showing preference for *stratum radiatum* GABAergic interneurons within the hippocampus (Li et al., 2021). For these reasons, we evaluated the ability of (+)-EU1180-453 to potentiate GluN1-1b-containing NMDARs.

In the presence of saturating co-agonists (1 mM glutamate and 100 µM glycine), 30 μM (+)-EU1180-453 potentiated GluN1-1b/GluN2C NMDARs to 202% of maximum response, with an EC_50_ of 0.6 μM (**Figure 3A,C**); GluN1-1b/GluN2D NMDARs were potentiated to 189% of maximum response, with an EC_50_ of 0.7 μM (**Figure 3B,D**). Using rapid solution-exchange (10 ms application) in transiently transfected HEK cells, GluN1-1b-containing NMDARs yielded similar results to those observed with the GluN1-1a splice variant. Overall charge transfer at both GluN1-1b/GluN2C and GluN1-1b/GluN2D NMDARs) were markedly and significantly increased after (+)-EU1180-453 application (**Figure 3E,F,G**). We also observed no instances of (+)-EU1180-453 induced agonist-dependency (via 10-90% rise time) or alteration of desensitization (via I_SS_/I_peak_ with 1.5 s agonist application time) (See **Supplemental Table S3**).

**Figure 3.**
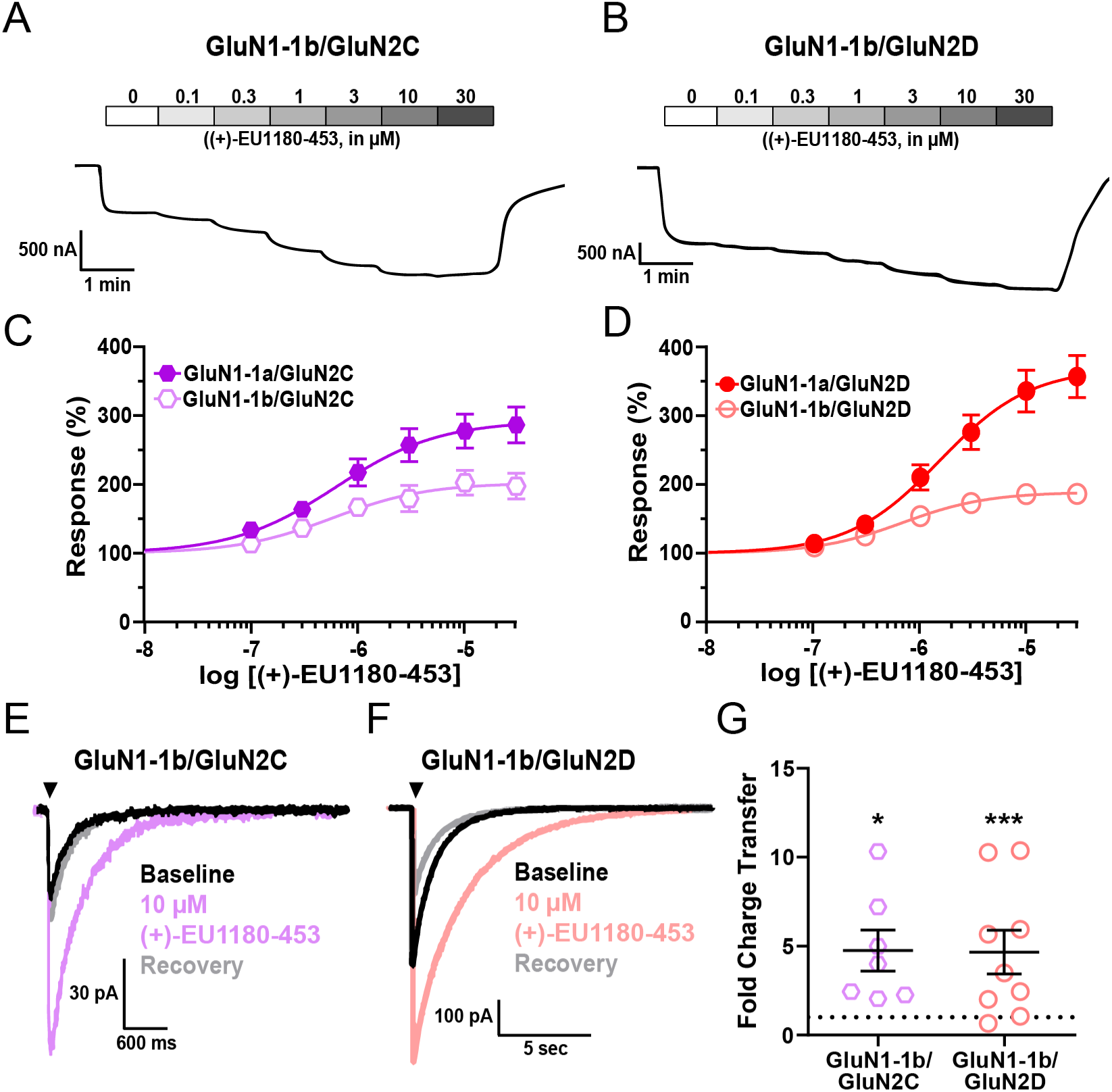
(+)-EU1180-453 potentiates response of exon5-containing (GluN1-1b) GluN2C and GluN2D NMDARs. **A)** Representative current responses from *Xenopus* oocytes of **A)** GluN1-1b/GluN2C and **B)** GluN1-1b/GluN2D NMDARs to varying concentrations of (+)-EU1180-453 in presence of saturating concentrations of glutamate (100 µM) and glycine (30 µM). Fitted concentration-response curves of **C)** GluN1-1b/GluN2C and **D)** GluN1-1b/GluN2D NMDARs to varying concentrations of (+)-EU1180-453 in presence of 30 µM glycine and 100 µM glutamate. GluN1-1a/GluN2C and GluN1-1a/GluN2D responses included as reference from Figure 2. Maximum potentiation of percent response (amplitude with saturating glycine and glutamate only) was **C)** 198 ± 19% of control for GluN1-1b/GluN2C and **D)** 187 ± 17% of control for GluN1-1b/GluN2D. Number of oocytes for each construct ranged from N=10-18. Representative responses from HEK293T cells are shown for baseline (glutamate only, black trace), glutamate plus 10 µM (+)-EU1180-453 (color trace), and glutamate recovery (gray trace) for **E)** GluN1-1b/GluN2C (n=7 cells) and **F)** GluN1-1b/GluN2D (n=9 cells). Cells were held at -70 mV and subjected to rapid and brief (10 ms) application of saturating glutamate (100 µM) in constant presence of 30 µM glycine. Current responses were recorded in absence, presence, and following removal of (+)-EU1180-453. **G)** The mean fold-change in charge transfer (response/baseline) is shown (paired t-test; baseline vs drug). The dotted line is at fold change of 1. All data are mean ± SEM. * p < 0.05; *** p < 0.001.

The overall level of potentiation of GluN1-1b-containing NMDARs with (+)-EU1180-453 was lower than those observed with GluN1-1a-containing NMDARs. Maximum potentiation at both GluN2C (202% for GluN1-1b vs 292% for GluN1-1a) and GluN2D (189% for GluN1-1b vs 370% for GluN1-1a) receptors was lower with exon5-containing receptors. The fold change in charge transfer was lower at GluN2C (4.8 ± 1.2 for GluN1-1b vs 6.5 ± 0.9 for GluN1-1a) receptors, but unchanged at GluN2D (4.7 ± 1.2 for GluN1-1b vs 4.7 ± 0.9 for GluN1-1a) receptors. These data indicate that the presence of exon5 within GluN1 may dampen, but does not inhibit, the ability of (+)-EU1180-453 to potentiate GluN2C- and GluN2D-containing NMDARs. Thus, NMDARs expressed on GABAergic interneurons within *stratum radiatum*, which are enriched for GluN2D-containing NMDARs (Perszyk et al., 2016; von Engelhardt et al., 2015), should be potentiated by (+)-EU1180-453.

### Stratum Radiatum GABAergic Interneurons, but Not CA1 Pyramidal Cells, are Potentiated by (+)-1180-453

(+)-1180-EU453 was assayed for activity at native, synaptic NMDARs in two different classes of neurons – CA1 pyramidal cells and *stratum radiatum* GABAergic interneurons. These two canonical neuronal classes were chosen as CA1 pyramidal cells primarily express GluN2A- and GluN2B-containing NMDARs (Hansen et al., 2014), whereas CA1 *stratum radiatum* GABAergic interneurons express functional GluN2A-, GluN2B-, and GluN2D-containing NMDARs (Perszyk et al., 2016; von Engelhardt et al., 2015). Hippocampal slices from juvenile (P11-15) WT mice were used to record NMDAR-mediated excitatory postsynaptic currents (EPSCs) onto CA1 pyramidal cells or CA1 *stratum radiatum* interneurons. Application of 10 µM (+)-EU1180-453 significantly enhanced charge transfer at CA1 *stratum radiatum* interneurons (**Figure 4F-H**; -6.2 ± 1.3 nA*ms for baseline vs -10.2 ± 2.0 nA*ms for (+)-EU1180-453), but was without effect on charge transfer at CA1 pyramidal cells (**Figure 4B-D**). Vehicle (0.1% DMSO) application did not alter charge transfer for either neuron class (**Figure 4B-H**). Additionally, there was no effect of (+)-EU1180-453 on rise time (**Supplemental Figure S4A**), peak amplitude (**Supplemental Figure S4B**) or weighted tau (**Supplemental Figure S4C**) at CA1 pyramidal cells. Both peak amplitude (-69 ± 11 pA for baseline vs -93 ± 15 pA for (+)-1180-453) and tau weighted (94 ± 15 ms for baseline vs 118 ± 18 ms for (+)-1180-453) were significantly increased following (+)-EU1180-453 application at CA1 *stratum radiatum* interneurons (**Supplemental Figure S4E,F**, **Supplemental Table S4)**. All measures described above were not influenced by changes in series resistance, which was monitored throughout recording in all cells (**Supplemental Figure S4G-J**).

**Figure 4.**
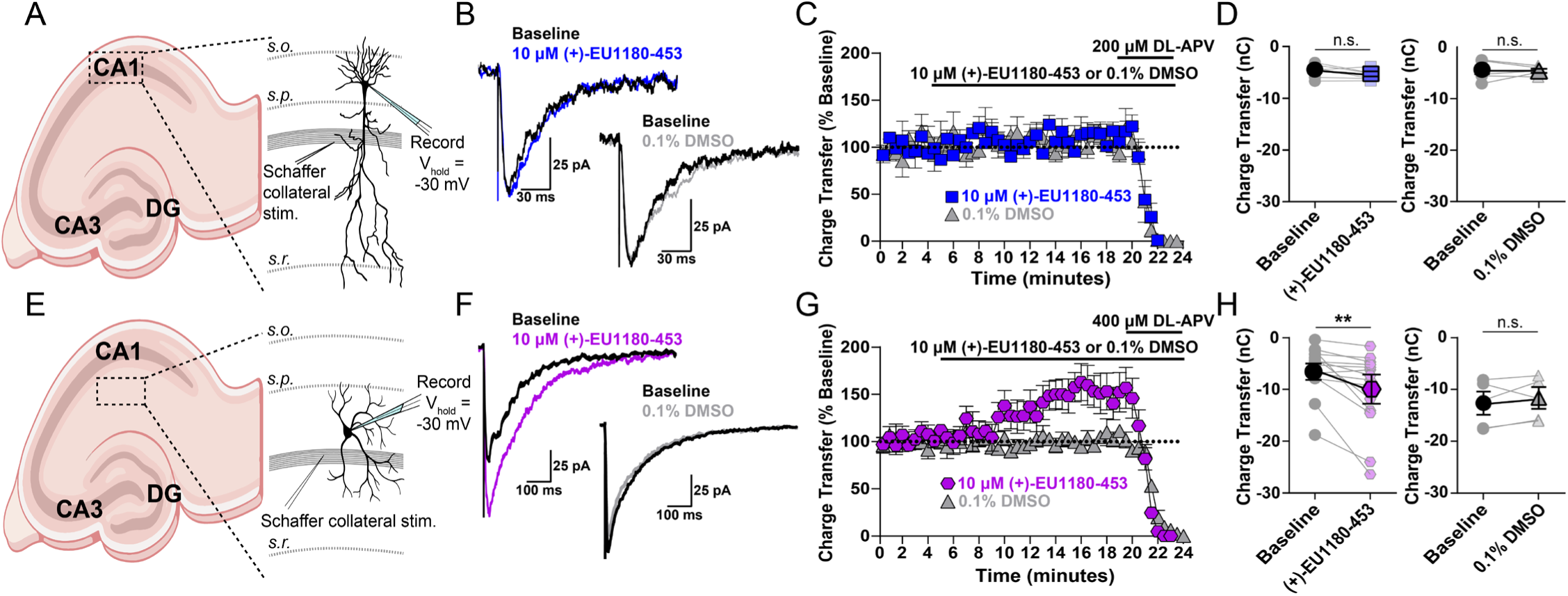
Synaptic NMDAR-mediated EPSCs in CA1 *stratum radiatum* GABAergic interneurons, but not CA1 pyramidal cells, are potentiated by (+)-EU1180-453 in developing hippocampus. CA1 *stratum radiatum* GABAergic interneurons or CA1 pyramidal cells were held at -30 mV in 0.2 mM extracellular Mg^2+^ and NMDAR-mediated EPSCs were pharmacologically isolated (see Methods). **A**) and **E**) Diagram of recording arrangement from mouse hippocampal slices. **B**) Representative EPSCs show the effects of 10 µM (+)-EU1180-453 (blue) or vehicle (gray) compared to baseline response (black) at CA1 pyramidal cells. **C**) Charge transfer vs time is shown for 10 µM (+)-EU1180-453 (blue) or vehicle (gray). Values are expressed as the percent of the average of 10 baseline responses. **D**) Application of 10 µM (+)-EU1180-453 (n = 6 cells, blue) or vehicle (n = 6 cells, 0.1% DMSO, gray) does not alter the charge transfer of NMDAR-mediated EPSCs onto CA1 pyramidal cells. **F**) Representative EPSCs show the effects of 10 µM (+)-EU1180-453 (magenta) or vehicle (gray) compared to baseline response (black) at *stratum radiatum* interneurons. **G**) Charge transfer vs time is plotted for 10 µM (+)-EU1180-453 (magenta) or vehicle (gray). Responses were calculated as a percent of the average of 10 baseline responses. **H**) Application of 10 µM (+)-EU1180-453 (n = 14 cells, magenta) potentiates the charge transfer of NMDAR-mediated EPSCs onto interneurons (-6.2 ± 1.3 nA*ms for baseline vs -10.2 ± 2.0 nA*ms for (+)-EU1180-453). Application of vehicle (n = 5 cells, 0.1% DMSO, gray) does not alter the charge transfer of NMDAR-mediated EPSCs onto interneurons. All data are mean ± SEM. Statistical significance determined by paired t-test. See **Supplemental Table S4** for full results. s.o. = *stratum oriens*; s.p. = *stratum pyramidale*; s.r. = *stratum radiatum*; CA1 = *cornu Ammonis* 1; CA3 = *cornu Ammonis* 3; DG = dentate gyrus; ** p < 0.01; n.s. indicates not significant.

Taken together, these data demonstrate the ability of (+)-EU1180-453 to selectively potentiate excitatory synaptic transmission onto hippocampal GABAergic interneurons. Despite (+)-EU1180-453 displaying very modest effects for GluN2A- and GluN2B-containing NMDARs in HEK cells, the lack of any observed potentiation at CA1 pyramidal cells suggests that these effects are below our limit of detection.

### (+)-EU1180-453 Enhances Inhibitory Tone in the Developing Hippocampal Circuit

Since (+)-1180-453 was able to potentiate the NMDAR-mediated EPSC onto CA1 *stratum radiatum* interneurons, we tested its ability to modulate the GABAergic interneuron network. For this, we recorded spontaneous inhibitory postsynaptic currents (sIPSCs) onto CA1 pyramidal cells from juvenile (P11-15) mouse hippocampal slices. We hypothesized that since (+)-1180-453 potentiates the excitatory drive onto inhibitory interneurons, (+)-1180-453 should enhance the output of these inhibitory cells onto their synaptic partners such as CA1 pyramidal cells. sIPSCs were recorded at V_hold_ +10 mV, the V_rev_ for ionotropic glutamate receptors. This eliminated the driving force for current flow through α-amino-3-hydroxy-5-methyl-4-isoxazolepropionic acid receptor (AMPAR) without the use of AMPAR antagonists, thus maintaining intact excitatory drive onto the interneuron network. Though this approach spares excitation within the slice, depolarizing pyramidal cells will generate depolarization-induced suppression of inhibition (DSI) via endocannabinoid release (Pitler and Alger, 1992; Wilson and Nicoll, 2001), which alters the overall content of sIPSCs recorded from various interneuron populations. More specifically, DSI preferentially inhibits GABA release from cholecystokinin-expressing basket cells but does not impact parvalbumin (PV)-expressing basket cells (Freund et al., 2003; Katona et al., 1999). Thus, the majority of our sIPSCs were likely mediated by PV basket cells, with minimal activity from CCK-expressing cells.

Application of 10 µM (+)-EU1180-453 showed a robust enhancement of inhibitory tone on CA1 pyramidal cells, significantly increasing the average sIPSC frequency (7.7 ± 0.9 Hz for baseline vs 10.2 ± 1.1 Hz for 10 µM (+)-EU1180-453) and significantly shifting the cumulative probability plot for inter-event interval to the left (**Figure 5**). There was no impact of vehicle on sIPSC frequency or inter-event interval (**Figure 5**). Application of 10 µM (+)-1180-453 showed no effect on average sIPSC amplitude (**Supplemental Figure S5A,E**). The cumulative probability plot for sIPSC amplitude was significantly shifted to the right after the application of (+)-EU1180-453 (**Supplemental Figure S5B**). There was no observed effect of vehicle on average sIPSC amplitude or cumulative probability event amplitude (**Supplemental Figure S5C-E**). We also observed no change in average sIPSC decay time for either (+)-EU1180-453 or vehicle (**Supplemental Figure S5F-J, Supplemental Table S5**) for more details. All changes observed to sIPSCs were not due to changes in series resistance, which was monitored at four separate times per cell during all recordings (**Supplemental Figure S5K-L**).

**Figure 5.**
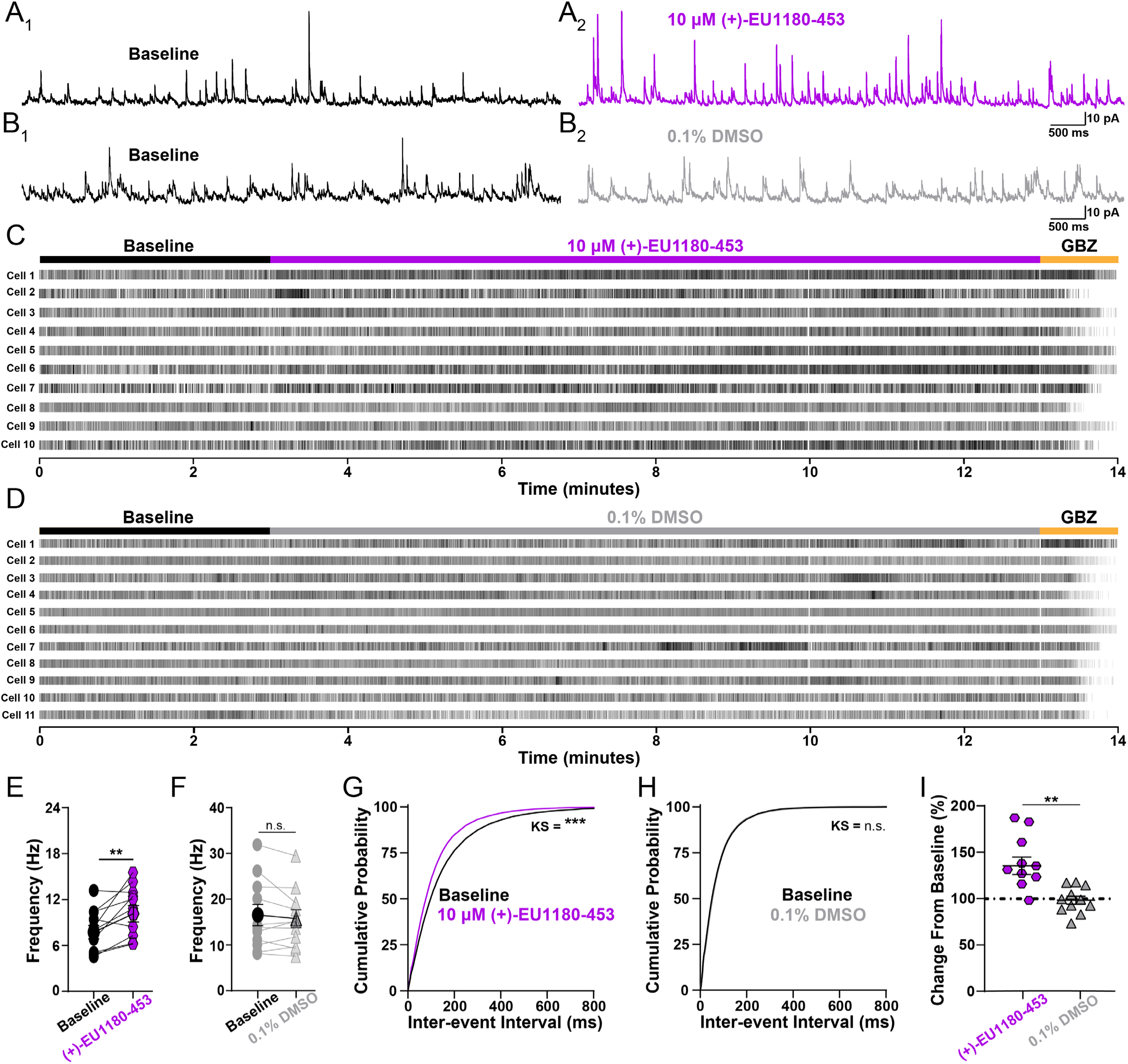
Application of (+)-EU1180-453 increases inhibitory tone in developing CA1. Spontaneous IPSCs (sIPSCs) were recorded at +10 mV from mouse CA1 pyramidal cells (P11-15). Representative sIPSCs **A_1_**) before (black) and **A_2_)** after 10 µM (+)-EU1180-453 (magenta, n = 11 cells), **B_1_**) before (black) and **B_2_**) after vehicle (0.1% DMSO, gray, n = 11 cells). Raster plots for all cells recorded in with **C**) (+)-EU1180-453 and **D**) 0.1% DMSO. **E**) Average sIPSC frequency is significantly increased following 10 µM (+)-EU1180-453 application (7.7 ± 0.9 Hz for baseline vs 10.2 ± 1.1 Hz for 10 µM (+)-EU1180-453; paired t-test), while **F**) there is no change in sIPSC frequency following 0.1% DMSO (paired t-test). **G**) Cumulative probability of interevent interval is significantly shifted to the left after 10 µM (+)-EU1180-453 application (KS test). **H**) Cumulative probability of interevent interval is unchanged in response 0.1% DMSO (KS test). **I**) Plots of average frequency percent change from baseline for 10 µM (+)-EU1180-453 (magenta) and (gray). 10 µM (+)-EU1180-453 application showed a significant increase the average frequency percent change from baseline compared to 0.1% DMSO (unpaired t-test). Data are mean ± SEM; GBZ = gabazine (SR-95531; 10 µM); KS = Kolmogorov-Smirnov; * p < 0.05; ** p < 0.01; *** p < 0.001; n.s. not significant.

To ensure the enhancement of sIPSC frequency we observed was indeed due to actions of (+)-EU1180-453 on GluN2D-containing NMDARs, we also recorded sIPSCs before and after modulator application onto CA1 pyramidal cells from juvenile (P11-15) homozygous *Grin2d* global knockout mice (*Grin2d*^-/-^). Application of 10 µM (+)-1180-453 significantly decreased sIPSC frequency (**Supplemental Figure S6A**); 2.1 ± 0.2 Hz for baseline vs 1.7 ± 0.2 Hz for 10 µM (+)-1180-453 in *Grin2d*^-/-^ mice. The cumulative probability plot for inter-event interval was significantly shifted to the right (**Supplemental Figure S6D**), opposite the shift in wild-type. Application of 10 µM (+)-1180-453 showed no effect on average sIPSC amplitude in *Grin2d*^-/-^ mice (**Supplemental Figure S6E**), however, the cumulative probability plot for sIPSC amplitude was significantly shifted to the left (**Supplemental Figure S6F**). These results support the idea that potentiation of GluN2D receptors on GABAergic interneurons can impact IPSC frequency onto pyramidal cells.

Taken together, these data demonstrate that potentiation of the NMDAR-mediated EPSC onto CA1 *stratum radiatum* interneurons is sufficient to drive an increase in inhibitory output onto excitatory pyramidal cells. This increase in GABAergic interneuron network activity is mediated by GluN2D-containing NMDARs, as we observed a *reduction* in sIPSC frequency with (+)-EU1180-453 when recording CA1 pyramidal cells from *Grin2d*^-/-^ mice. Although this result in *Grin2d*^-/-^ mice was surprising, it could stem from altered expression of NMDARs, perhaps with GluN2C-containing NMDARs being expressed presynaptically. Alternatively, astrocytes also express GluN2C NMDARs in WT mice (Chipman et al., 2021; Shelkar et al., 2021) and modulation of astrocytic GluN2C-containing NMDARs may impact inhibitory tone via modulation of glutamate and/or GABA uptake. Regardless, these data highlight a clear and robust utility for GluN2D modulation in the role of network inhibition and interneuron output.

### (+)-EU1180-453 Alters Excitatory-Inhibitory Balance and Enhances Carbachol-Induced Gamma Band Power in the Developing Hippocampal Circuit

To end, we explored how (+)-EU1180-453 influences excitatory-inhibitory (E-I) tone and network activity. Given the specificity of (+)-EU1180-453 to modulate the interneuron network, we hypothesized (+)-EU1180-453 would also shift excitatory-inhibitory balance. We recorded compound excitatory postsynaptic potentials (EPSPs) and inhibitory postsynaptic potentials (IPSPs) in developing CA1 pyramidal cells. Stimulation of Schaffer collaterals cause CA3 afferents to generate excitatory glutamatergic signaling in both CA1 pyramidal cells and CA1 interneurons. This then causes CA1 interneurons to fire, generating an IPSP shortly after the CA3-mediated EPSP as recorded from CA1 pyramidal cells. We used this paradigm to assess E-I coupling in CA1 pyramidal cells, an important subset projection neuron. Application of 10 µM (+)-EU1180-453 caused a significant shift in the E-I ratio, while vehicle controls showed no change (**Figure 6A-C**). We also observed a significant increase in IPSP amplitude as well as significant decrease in EPSP amplitude after 10 µM (+)-EU1180-453 (**Supplemental Figure S7A**,**B**; **Supplemental Table S6**). Taken together, these data demonstrate that modulation of NMDARs on interneurons only is capable of altering E-I coupling.

**Figure 6.**
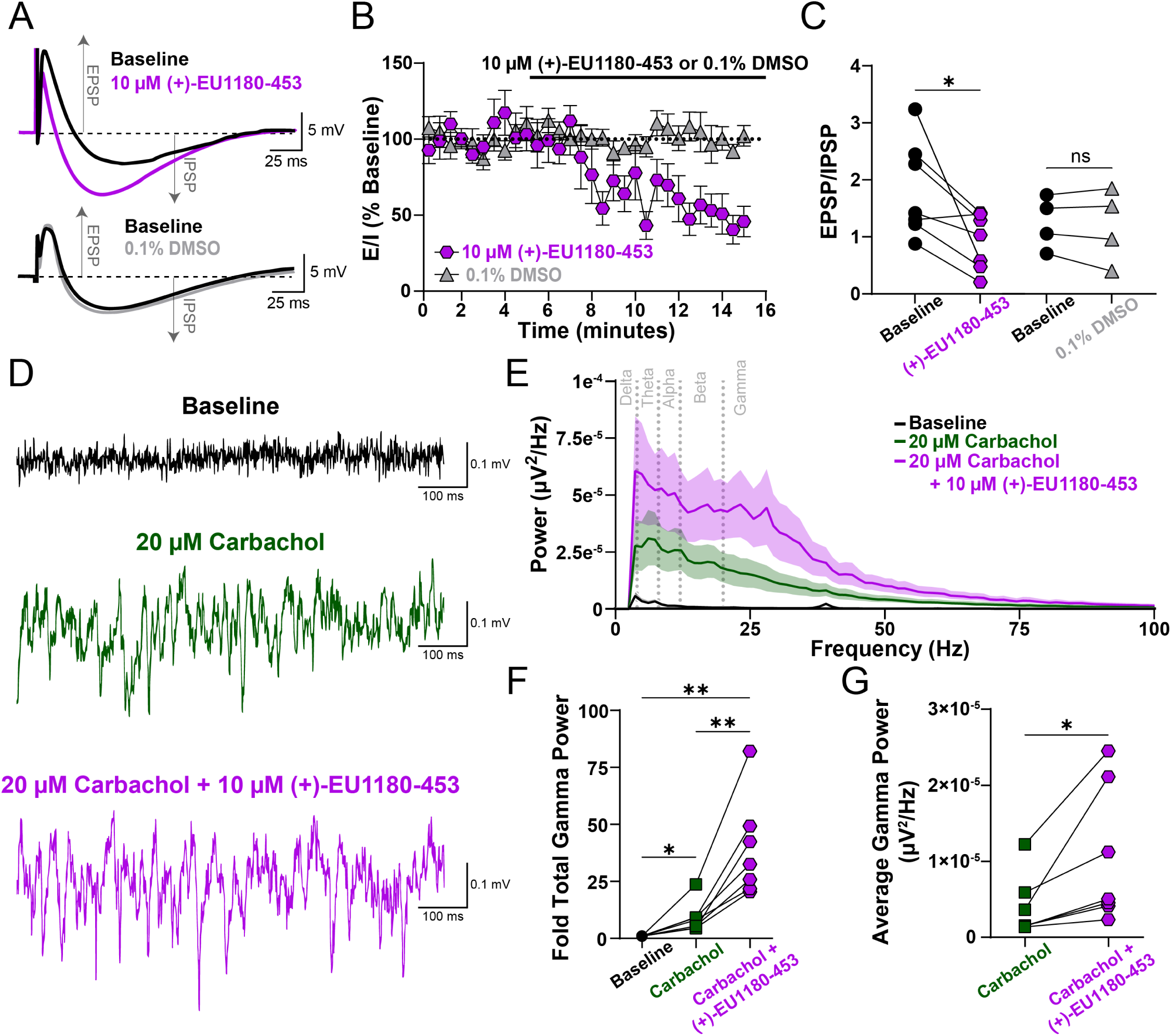
(+)-EU1180-453 shifts excitatory-inhibitory ratio towards inhibition and enhances carbachol-induced gamma power in CA1 of hippocampus. **A**) Representative traces of dual excitatory postsynaptic potential (EPSP) and inhibitory postsynaptic potential (IPSP) before (black) and after 10 µM (+)-EU1180-453 (magenta) or before (black) and after 0.1% DMSO (gray). **B**) Time series plot showing EPSP/IPSP ratio before and after wash-in of either 10 µM (+)-EU1180-453 (magenta) or 0.1% DMSO (gray). **C**) EPSP/IPSP ratio is significantly reduced after (+)-EU1180-453 application (2.0 ± 0.3 baseline vs 0.9 ± 0.2 (+)-EU1180-453; paired t-test) but unchanged with 0.1% DMSO. **D**) Representative field potential recordings from CA1 *stratum pyramidale* at baseline (black), after 20 µM carbachol (green), and after 20 µM carbachol plus 10 µM (+)-EU1180-453 (magenta). **E**) Average power spectral density plot at baseline (black), after 20 µM carbachol (green), and after 20 µM carbachol plus 10 µM (+)-EU1180-453 (magenta). Dark line is the average power, with shaded areas representing the SEM. **F**) Fold change in total gamma band power compared to baseline for 20 µM carbachol only and 20 µM carbachol plus 10 µM (+)-EU1180-453. There are significant increases in fold gamma band power after 20 µM carbachol only, between 20 µM carbachol only and 20 µM carbachol plus 10 µM (+)-EU1180-453, and baseline and 20 µM carbachol plus 10 µM (+)-EU1180-453 (one-way ANOVA). **G**) Carbachol-induced average gamma band power is significantly increased by application 10 µM (+)-EU1180-453 (paired t-test). Data are mean ± SEM. EPSP = excitatory postsynaptic potential; IPSP = inhibitory postsynaptic potential; * = p < 0.05; ** = p < 0.01; **** = p < 0.0001; ns = not significant.

To explore the impact of (+)-EU1180-453 on neural oscillations, we used 20 µM carbachol to elevate hippocampal slice excitability *in vitro* and recorded extracellular field potentials in *stratum pyramidale* in developing hippocampus. More specifically, we used this approach to simulate neural activity of an awake, behaving animal as exogenous carbachol application to an *in vitro* brain slice has been shown to generate gamma oscillations via field potential recordings (Fellous and Sejnowski, 2000). Gamma band oscillations *in vivo* have been recorded from both rodents (Traub et al., 1996) and humans (Le Van Quyen et al., 2010) and are thought to represent neural computation associated with memory recall and stimulus discrimination (Herrmann et al., 2004). More specifically, gamma oscillations are largely driven by perisomatic inhibition from fast-spiking parvalbumin-positive interneurons (Buzsaki and Wang, 2012; Cardin et al., 2009; Carlen et al., 2012) and have been shown to be disrupted in humans with various forms of neuropsychiatric disorders (Fitzgerald and Watson, 2018; Light et al., 2006).

Application of 10 µM (+)-EU1180-453 potentiated carbachol-induced gamma band field oscillations. Compared to baseline, (+)-EU1180-453 increased the fold change in total gamma band power by 31-fold compared to only 14-fold with carbachol alone (**Figure 6E-F**). Average gamma band power was also significantly increased when compared to carbachol alone (39 ± 15 µV^2^/Hz *10^7^ for carbachol alone vs 104 ± 34 µV^2^/Hz *10^7^ for carbachol plus (+)-EU1180-453; **Figure 6G**). We also observed significant increases in baseline-normalized total power in the alpha (8-12 Hz) and beta (12-20 Hz) when comparing carbachol alone to carbachol plus (+)-EU-1180-453 (**Supplemental Figure S7F**; **Supplemental Table S6**). Together, these data highlight that modulation of excitatory NMDAR-mediated signaling in GABAergic interneurons is sufficient to alter E-I coupling in projection neurons as well as modulation neural network oscillations. Increasing the power output of gamma oscillations *in vitro* is promising data to suggest that these oscillations could be modified *in vivo* in preclinical rodent models of neuropsychiatric disease.

## Discussion

NMDAR modulation has potential therapeutic benefit for multiple indications. Here, we describe a novel GluN2C/GluN2D PAM, (+)-EU1180-453, which has improved drug-like properties over the previous first-in-class GluN2C/GluN2D-selective PAM (+)-CIQ (Epplin et al., 2020). (+)-EU1180-453 possesses increased aqueous solubility and increased doubling concentration compared to the prototypical GluN2C/GluN2D PAM (+)-CIQ, without compromising drug specificity and efficacy. (+)-EU1180-453 potentiates triheteromeric NMDARs containing at least one GluN2C or GluN2D subunit and is active at both major splice variants of the GluN1 subunit (GluN1-1a and GluN1-1b). Heterologous HEK cell recordings reveal (+)-EU1180-453 potentiation is not agonist-dependent and increases receptor efficacy by both prolonging deactivation time and potentiating peak amplitude. Using developing mouse hippocampal slices, we show that (+)-EU1180-453 selectively increases synaptic NMDAR-mediated charge transfer onto CA1 *stratum radiatum* interneurons while having no detectable effect on synaptic NMDARs expressed on pyramidal cells. This increase in NMDAR charge transfer is sufficient to generate a significant increase in inhibitory output from GABAergic interneurons that is dependent on the GluN2D subunit. (+)-EU1180-453 also shifts excitatory-to-inhibitory coupling towards increased inhibition and produces enhanced gamma band power from carbachol-induced field potential oscillations in hippocampus. By increasing the overall excitatory tone onto GABAergic interneurons, (+)-EU1180-453 enhances circuit inhibition, which could prove advantageous for numerous neuropsychiatric diseases such as anxiety, depression, epilepsy, and schizophrenia.

These data highlight the utility of (+)-EU1180-453 as a tool compound for future *in vitro* and *in vivo* experiments and also raise important questions about NMDAR modulation and neurophysiology. Triheteromeric NMDARs are believed to comprise a significant proportion of NMDAR-mediated synaptic current (Hansen et al., 2014; Kim et al., 2005; Stroebel et al., 2018), however, the exact ratio of triheteromeric and diheteromeric NMDARs at native synapses remains elusive (Goncalves et al., 2020; Kellermayer et al., 2018). Moreover, the repertoire of possible triheteromeric NMDAR combinations compared to those expressed in native tissue is also unknown. For example, we did not test (+)-EU1180-453 on potential GluN2B/GluN2C and GluN2C/GluN2D triheteromeric complexes, which have not been extensively studied. We also did not explore the possibility of (+)-EU1180-453 potentiation of triheteromeric complexes containing GluN2C or GluN2D with GluN3A or GluN3B. Our previous data indicates (+)-EU1180-453 does not potentiate GluN3A diheteromeric NMDARs and shows slight inhibition of GluN3B diheteromeric NMDARs (Epplin et al., 2020). Our current understanding of potential triheteromeric NMDARs complexes with one GluN2 subunit and GluN3 subunit is incomplete (Pachernegg et al., 2012).

In addition to a multitude of different GluN2 or even GluN3 subunit combinations, there are also eight different splice variants of the obligate GluN1 subunit (Hansen et al., 2021). Canonically, these GluN1 splice variants have been stratified as either containing or lacking 21 amino acids in the amino-terminal domain encoded by an alternatively spliced exon. The bulk of heterologous studies have been conducted on NMDARs that’s lack the 21 amino acid aminoterminal domain insert, as this splice variant is believed to be predominantly expressed in native tissue (Laurie and Seeburg, 1994). Nonetheless, inclusion of these 21 residues in the amino terminal domain alters several channel properties. That is, inclusion of this alternative exon accelerates the deactivation time course (Rumbaugh et al., 2000; Vance et al., 2012), alters channel desensitization (Rumbaugh et al., 2000), reduces proton and polyamine sensitivity (Traynelis et al., 1995), reduces modulation by extracellular zinc ions (Traynelis et al., 1998), and endows non-ionotropic properties such as glycine priming (Li et al., 2021). The distribution of the GluN1 amino terminal domain splice variant displays a cell-type specific expression profile, with most excitatory pyramidal cells expressing GluN1-1a variants (exon5-lacking) and *stratum radiatum* GABAergic interneurons expressing the GluN1-1b variant (exon5-containing) (Li et al., 2021). Although we report that (+)-EU1180-453 is active at both GluN1-1a and GluN1-1b splice variants, activity for GluN1-1b is lower than GluN1-1a. This lowered potentiation may help to explain why we only observed a ∼50% increase in evoked NMDAR-mediated EPSCs onto *stratum radiatum* interneurons.

An alternative explanation for the lower-than-expected potentiation of (+)-EU1180-453 at NMDARs on *stratum radiatum* interneurons may be due to the heterogeneity of the interneuron population within the hippocampus. Within the CA1 region alone, there are more than 15 different subtypes of GABAergic interneurons (Pelkey et al., 2017). Focusing on just the *stratum radiatum* cell layer within CA1, there are at least six different types of interneurons, with the majority being from the caudal ganglionic eminence (CGE) (Pelkey et al., 2017). The majority of *stratum radiatum* interneurons are cholecystokinin (CCK)-expressing interneurons as well as Ivy cells and some vasoactive intestinal peptide (VIP)-expressing cells. Within the CCK interneuron class, there are five different subtypes within the *stratum radiatum*: VGluT3-expressing basket cells, VIP-expressing basket cells, Schaffer collateral-associated cells, apical dendrite-innervating cells, and perforant path-associated cells. Thus, the GABAergic interneuron landscape within *stratum radiatum* is vast and different subtypes of interneuron may express different combinations of NMDAR subunits. While it is appreciated that CCK-expressing interneurons express the GluN2D subunit, there has yet to be a comprehensive evaluation of NMDAR subtype expression across interneuron subgroups. Additionally, it is currently unknown whether Ivy cells and VIP cells express GluN2D. NMDAR subunit expression is also activity-dependent, and studied best in the context of the GluN2B-to-GluN2A switch (Carmignoto and Vicini, 1992), although activity-dependency likely modulates expression of other NMDAR subunits as well.

Our measurement of interneuron output – frequency of sIPSCs onto CA1 pyramidal cells – was obtained by depolarizing pyramidal cells to the reversal potential for ionotropic glutamate receptors (between 0 mV and +10 mV). In doing this, we were able to preserve excitatory tone onto GABAergic interneurons, however, depolarization of pyramidal cells induces DSI (Pitler and Alger, 1992; Wilson and Nicoll, 2001), which preferentially inhibits GABA release from CCK-expressing basket cells, diminishing their contribution to sIPSCs onto pyramidal cells (Freund et al., 2003; Katona et al., 1999). Thus, we were able to show robust enhancement of interneuron output with only a fraction of the total interneurons within the circuit functioning. Knockout studies combined with NMDAR subunit-selective pharmacological tools suggests that parvalbumin-expressing basket cells express GluN2A, GluN2B, and GluN2D NMDARs (Booker et al., 2021; Gawande et al., 2023; Hanson et al., 2019) while CCK-expressing basket cells only express GluN2B and GluN2D subunits (Booker et al., 2021).

When evaluating our E-I coupling and carbachol-induced gamma oscillations in field potential recordings, our data comprise the entire interneuron network. Schaffer collateral afferents innervate all CA1 interneurons (Pelkey et al., 2017) and thus each unique class of GABAergic interneuron expressed within CA1 could contribute to our E-I coupling data. Given that we used monopolar stimulation electrodes in the upper third of *stratum radiatum*, however, it is more likely that we are generating IPSPs from PV and CCK basket cells, with some contributions from other interneurons classes. Carbachol provides broad excitatory drive to neurons throughout hippocampus, increasing both excitatory and inhibitory synaptic activity, spike firing, and thus fluctuations in extracellular field potential recordings. Carbachol-evoked gamma oscillations are shown to be largely driven by PV-expressing interneurons (Buzsaki and Wang, 2012; Cardin et al., 2009; Carlen et al., 2012), with some contributions from somatostatin (SST)-expressing interneurons (Antonoudiou et al., 2020). Moreover, synaptic GluN2D expression has been demonstrated for cortical PV cells (Gawande et al., 2023) and *Grin2d* mRNA has been reported in cortical SST cells (Huntley et al., 2020; Perszyk et al., 2016).

It is important to note that (+)-EU1180-453 may also be impacting astrocyte function. Hippocampal astrocytes have robust expression of the GluN2C subunit (Chipman et al., 2021), with recent data from knockout studies suggesting that astrocytes express GluN2A/GluN2C triheteromeric NMDARs (Shelkar et al., 2022). Given that (+)-EU1180-453 can potentiate GluN2C-containing NMDARs and GluN2A/GluN2C triheteromeric receptors, it seems probable that our compound could impact astrocyte function. The potential role of astrocytic involvement in interneuron output is currently unknown.

These data highlight that changing the excitatory tone onto GABAergic interneurons is sufficient to increase overall inhibitory output within a circuit. While E-I alteration hypotheses in the neuropsychiatric field have been proposed for decades, data demonstrating that the interneuron network *can be* enhanced has been lacking. This manuscript provides proof-of-concept that interneuron modulation may be a tractable drug target for disease. Interneuron dysfunction is a known phenomenon in the diseased brain and has been causally linked to etiology of numerous neuropsychiatric disorders. Now, with this improved tool compound, preclinical rodent models of neuropsychiatric disease can be tested for efficacy of interneuron elevation in relief of aberrant circuit function. If successful, targeted interneuron genetic therapies aimed at increasing inhibitory output could prove beneficial to symptom modulation.

## Ethics Statement

### Conflict of interest

D.C.L., H.Y., M.P.E., N.S.A., and S.F.T. are co-inventors of Emory-owned intellectual property. S.F.T. is a member of the SAB for Sage Therapeutics, Eumentis Therapeutics, Neurocrine, the GRIN2B Foundation, the CureGRIN Foundation, and CombinedBrain. S.F.T. is a consultant for GRIN Therapeutics. H.Y. is the PI on a research grant from Sage Therapeutics and GRIN Therapeutics to Emory. S.F.T. is PI on a research grant from GRIN Therapeutics to Emory. S.F.T. is cofounder of NeurOp, Inc. and Agrithera. D.C.L. and S.F.T. are on the Board of Directors for NeurOp. T.A.B. is a member of the SAB for GRIN2B Foundation, CureGRIN Foundation and GRIN Therapeutics; all remuneration has been made to his department.

### Grant Support

CRC received grant support from NINDS (NS113530). This work was supported by a grant from the Uplifting Athletes Young Investigator Draft (REP). HY was supported by NIA (AG075444, AG079956 and AG081401) and NICHD (HD082373). SFT received grant support from NINDS (NS111619). This work was also supported by research grants to Emory University from Biogen and Janssen Research and Development (SFT).

## Abbreviations

AMPA: α-amino-3-hydroxy-5-methyl-4-isoxazolepropionic acid
APV: 2-amino-5-phosphonopentanoic acid
ATD: amino-terminal domain
CCK: cholecytsokinin
CIQ: (3-chlorophenyl)(6,7-dimethoxy-1-((4-methoxyphenoxy)-methyl)-3,4-dihydroisoquinolin-2(1*H*)-yl)methanone)
CNS: central nervous system
DMSO: dimethyl sulfoxide
DSI: depolarization-induced suppression of inhibition
EPSC: excitatory postsynaptic current
ER: endoplasmic reticulum
GABA: gamma-aminobutryic acid
NAM: negative allosteric modulator
NMDAR: N-methyl-D-aspartate receptor
P: postnatal day
PAM: positive allosteric modulator
PV: parvalbumin
sIPSC: spontaneous inhibitory postsynaptic current
VIP: vasoactive intestinal peptide.

## Acknowledgements

We thank Sukhan Kim for excellent technical assistance. We would also like to thank helpful conversations with Ken Pelkey and Chris McBain which sparked experimental design and data interpretation.

## Author Contributions

*Participated in research design*: Camp, Perszyk, Yuan, and Traynelis

*Conducted experiments*: Camp, Banke, Xing, Yu, and Zhang

*Contributed new reagents*: Epplin and Liotta

*Performed data analysis*: Camp, Banke, Perszyk, and Traynelis

*Contributed to writing of the manuscript*: Camp, Banke, Xing, Yu, Perszyk, Epplin, Zhang, Benke, Yuan, Liotta, and Traynelis

