## Supplemental Data and Tables for "Selective Enhancement of the Interneuron Network and Gamma-Band Power via GluN2C/GluN2D NMDA Receptor Potentiation"

| | $EC_{50}$, µM (95% CI) | Max Potentiation, % (± SEM) | n (cells) |
|---|---|---|---|
| *Diheteromeric NMDARs with C1- and C2-tails* | | | |
| **GluN1/GluN2A$_{C1}$/GluN2A$_{C2}$** | 8.4 (1.9-15) | 108 ± 0.8 | 12 |
| **GluN1/GluN2B$_{C1}$/GluN2B$_{C2}$** | 19 (13-24) | 124 ± 2.0 | 12 |
| **GluN1/GluN2C$_{C1}$/GluN2C$_{C2}$** | 0.8 (0.6-1.1) | 287 ± 26 | 12 |
| **GluN1/GluN2D$_{C1}$/GluN2D$_{C2}$** | 1.7 (1.5-1.9) | 367 ± 9.9 | 18 |
| *Triheteromeric NMDARs* | | | |
| **GluN1/GluN2A$_{C1}$/GluN2B$_{C2}$** | 2.7 (0.2-5.5) | 103 ± 2.2 | 12 |
| **GluN1/GluN2A$_{C1}$/GluN2C$_{C2}$** | 1.6 (1.0-2.1) | 141 ± 13 | 12 |
| **GluN1/GluN2A$_{C1}$/GluN2D$_{C2}$** | 1.8 (0.6-2.9) | 153 ± 16 | 15 |
| **GluN1/GluN2B$_{C1}$/GluN2D$_{C2}$** | 1.7 (1.3-2.0) | 183 ± 5.6 | 14 |
| *Exon5-Containing NMDARs* | | | |
| **GluN1-1b/GluN2C** | 0.5 (0.2-0.7) | 197 ± 19 | 14 |
| **GluN1-1b/GluN2D** | 1.0 (0.7-1.3) | 187 ± 12 | 12 |

**Supplemental Table S1**. Concentration-response relationships for (+)-EU1180-453 at various NMDAR subunits recorded from *Xenopus* oocytes. All recordings performed in presence of saturating concentrations of co-agonists (100 µM glutamate and 30 µM glycine). Maximum potentiation values reported as percent steady-state current with 30 µM (+)-EU1180-453 to steady-state current with 0 µM (+)-EU1180-453. N is number of oocytes. Mean and CI determined from the Log of the EC50.

| GluN2A *(n = 7 cells)* | $I_{peak}$ (pA) | Rise Time (ms) | Deactivation $T_W$ (ms) | Charge Transfer (nA*ms) | Fold Charge Transfer |
|---|---|---|---|---|---|
| **Control** | -1,700 ± 290 | 11 ± 0.9 | 41.0 ± 3.8 | -2.7 ± 0.7 | 1.0 ± 0 |
| **10 µM (+)-EU1180-453** | -1,900 ± 414 | 13 ± 1.2 | 63.0 ± 9.7 | -4.8 ± 1.0* | 1.9 ± 0.3 |
| **Recovery** | -1,520 ± 376 | 12 ± 0.7 | 38.0 ± 4.0 | -2.0 ± 0.4 | 0.8 ± 0.1 |

| GluN2B *(n =9 cells)* | $I_{peak}$ (pA) | Rise Time (ms) | Deactivation $T_W$ (ms) | Charge Transfer (nA*ms) | Normalized Charge Transfer |
|---|---|---|---|---|---|
| **Control** | -317 ± 90 | 19 ± 0.5 | 676 ± 62 | -8.8 ± 2.7 | 1.0 ± 0 |
| **10 µM (+)-EU1180-453** | -365 ± 100 | 21 ± 1.1 | 804 ± 93 | -12 ± 3.8 | 1.4 ± 0.1 |
| **Recovery** | -237 ± 62 | 20 ± 0.8 | 654 ± 91 | -6.0 ± 1.9 | 0.7 ± 0.1 |

| GluN2C *(n = 7 cells)* | $I_{peak}$ (pA) | Rise Time (ms) | Deactivation $T_W$ (ms) | Charge Transfer (nA*ms) | Normalized Charge Transfer |
|---|---|---|---|---|---|
| **Control** | -80.0 ± 30 | 26 ± 1.3 | 608 ± 79 | -1.4 ± 0.4 | 1.0 ± 0 |
| **10 µM (+)-EU1180-453** | -181 ± 48 | 24 ± 1.3 | 1,120 ± 86 | -9.9 ± 4.0* | 6.5 ± 0.9 |
| **Recovery** | -74.0 ± 32 | 28 ± 1.2 | 597 ± 63 | -1.4 ± 0.6 | 0.9 ± 0.1 |

| GluN2D *(n = 8 cells)* | $I_{peak}$ (pA) | Rise Time (ms) | Deactivation $T_W$ (ms) | Charge Transfer (nA*ms) | Normalized Charge Transfer |
|---|---|---|---|---|---|
| **Control** | -136 ± 32 | 28 ± 2.6 | 5,950 ± 481 | -31 ± 9.8 | 1.0 ± 0 |
| **10 µM (+)-EU1180-453** | -275 ± 48 | 22 ± 1.9 | 11,100 ± 609* | -110 ± 21* | 4.7 ± 0.9 |
| **Recovery** | -88.0 ± 19 | 27 ± 1.7 | 5,470 ± 712 | -19 ± 5.6 | 0.6 ± 0.1 |

*Exon5-Containing NMDARs*

| GluN1-1b/GluN2C *(n = 7 cells)* | $I_{peak}$ (pA) | Rise Time (ms) | Deactivation $T_W$ (ms) | Charge Transfer (nA*ms) | Normalized Charge Transfer |
|---|---|---|---|---|---|
| **Control** | -300 ± 136 | 8.1 ± 2.5 | 304 ± 31 | -89 ± 43 | 1.0 ± 0 |
| **10 µM (+)-EU1180-453** | -500 ± 243 | 8.1 ± 1.5 | 859 ± 131 | -44 ± 210* | 4.8 ± 1.1 |
| **Recovery** | -213 ± 83 | 9.2 ± 1.6 | 401 ± 52 | -84 ± 33 | 1.2 ± 0.2 |

| GluN1-1b/GluN2D *(n = 9 cells)* | $I_{peak}$ (pA) | Rise Time (ms) | Deactivation $T_W$ (ms) | Charge Transfer (nA*ms) | Normalized Charge Transfer |
|---|---|---|---|---|---|
| **Control** | -498 ± 217 | 5.2 ± 0.7 | 1320 ± 76 | -644 ± 340 | 1.0 ± 0 |
| **10 µM (+)-EU1180-453** | -610 ± 145 | 5.5 ± 1.7 | 2800 ± 149* | -15000 ± 370* | 4.7 ± 1.2 |
| **Recovery** | -320 ± 91 | 5.1 ± 0.7 | 1330 ± 92 | -390 ± 130 | 1.1 ± 0.3 |

**Supplemental Table S2**. NMDAR responses in transfected HEK293T cells exposed to brief duration 1 mM glutamate (control and recovery) or 1 mM glutamate plus 10 µM (+)-EU1180-453 for 10 ms in constant presence of 100 µM glycine. All recordings completed at $V_{hold}$ = -70 mV. Control (glutamate), drug application ((+)-EU1180-453 plus glutamate and glycine), and recovery (glutamate only) were repeated in the same cell. Deactivation Tw is the weighted deactivation time. Fold charge transfer is fold change of charge transfer in the drug application and recovery periods compared to the baseline period. N is the number of HEK293T cells. * = statistically significant after post-hoc correction of p-value for family-wise error; all statistical tests are paired t-tests compared to baseline.

| GluN2A<br>*(n = 8 cells)* | $I_{peak}$ (pA) | $I_{SS}$ (pA) | $I_{SS}$ / $I_{peak}$ (%) | Deactivation $T_W$ (ms) | Charge Transfer (nA*ms) |
|---|---|---|---|---|---|
| **Control** | -1,690 ± 236 | -1,140 ± 184 | 69 ± 6.2 | 79 ± 19 | -1,840 ± 323 |
| **10 µM (+)-EU1180-453** | -1,900 ± 344 | -1,050 ± 129 | 63 ± 7.6 | 107 ± 29 | -2,010 ± 300 |
| **Recovery** | -1,550 ± 343 | -604 ± 93 | 53 ± 10 | 78 ± 25 | -1,350 ± 241 |

| GluN2B<br>*(n = 9 cells)* | $I_{peak}$ (pA) | $I_{SS}$ (pA) | $I_{SS}$ / $I_{peak}$ (%) | Deactivation $T_W$ (ms) | Charge Transfer (nA*ms) |
|---|---|---|---|---|---|
| **Control** | -346 ± 97 | -309 ± 85 | 90 ± 0.7 | 899 ± 77 | -12.5 ± 3.6 |
| **10 µM (+)-EU1180-453** | -391 ± 103 | -301 ± 65 | 82 ± 4.4 | 1,150 ± 118 | -19.5 ± 6.1* |
| **Recovery** | -265 ± 68 | -173 ± 38 | 72 ± 7.0 | 1,040 ± 169 | -12 ± 4.0 |

| GluN2C<br>*(n = 7 cells)* | $I_{peak}$ (pA) | $I_{SS}$ (pA) | $I_{SS}$ / $I_{peak}$ (%) | Deactivation $T_W$ (ms) | Charge Transfer (nA*ms) |
|---|---|---|---|---|---|
| **Control** | -140 ± 63 | -140 ± 61 | 102 ± 1.3 | 692 ± 30 | -3.3 ± 1.4 |
| **10 µM (+)-EU1180-453** | -185 ± 35 | -188 ± 35 | 102 ± 0.9 | 1,570 ± 155* | -11 ± 3.1* |
| **Recovery** | -83 ± 27 | -85 ± 27 | 103 ± 2.0 | 881 ± 141 | -2.6 ± 1.0 |

| GluN2D<br>*(n = 8 cells)* | $I_{peak}$ (pA) | $I_{SS}$ (pA) | $I_{SS}$ / $I_{peak}$ (%) | Deactivation $T_W$ (ms) | Charge Transfer (nA*ms) |
|---|---|---|---|---|---|
| **Control** | -186 ± 45 | -189 ± 46 | 101 ± 1.0 | 6,420 ± 522 | -44 ± 14 |
| **10 µM (+)-EU1180-453** | -350 ± 51 | -338 ± 49 | 97 ± 0.7 | 13,200 ± 791* | -172 ± 35* |
| **Recovery** | -127 ± 23 | -124 ± 24 | 98 ± 2.4 | 7,310 ± 605 | -35 ± 8.3 |

*Exon5-Containing NMDARs*

| GluN1-1b/GluN2C<br>*(n = 7 cells)* | $I_{peak}$ (pA) | $I_{SS}$ (pA) | $I_{SS}$ / $I_{peak}$ (%) | Deactivation $T_W$ (ms) | Charge Transfer (nA*ms) |
|---|---|---|---|---|---|
| **Control** | -369 ± 168 | -320 ± 164 | 79 ± 3.7 | 439 ± 67 | -0.61 ± 0.3 |
| **10 µM (+)-EU1180-453** | -536 ± 237 | -490 ± 226 | 87 ± 2.7 | 1050 ± 286 | -1.23 ± 0.5* |
| **Recovery** | -278 ± 98 | -246 ± 94 | 82 ± 4.6 | 648 ± 155 | -0.52 ± 0.2 |

| GluN1-1b/GluN2D<br>*(n = 9 cells)* | $I_{peak}$ (pA) | $I_{SS}$ (pA) | $I_{SS}$ / $I_{peak}$ (%) | Deactivation $T_W$ (ms) | Charge Transfer (nA*ms) |
|---|---|---|---|---|---|
| **Control** | -604 ± 243 | -525 ± 223 | 83 ± 2.3 | 1620 ± 100 | -1.56 ± 0.6 |
| **10 µM (+)-EU1180-453** | -649 ± 136 | -576 ± 129 | 86 ± 2.2 | 2930 ± 142* | -2.44 ± 0.5* |
| **Recovery** | -441 ± 123 | -394 ± 119 | 85 ± 2.4 | 1920 ± 256 | -1.17 ± 0.3 |

**Supplemental Table S3**. NMDAR responses in transfected HEK293T cells exposed to prolonged application of 1 mM glutamate (control and recovery) or 1 mM glutamate plus 10 µM (+)-EU1180-453 for 1.5 seconds. All recordings completed in presence of 100 µM glycine and held at $V_{hold}$ = -70 mV. Control (glutamate and glycine only), drug application ((+)-EU1180-453 plus glutamate and glycine), and recovery (back to glutamate and glycine only) were repeated in the same cell. $I_{ss}$/$I_{peak}$ is percent of steady-state current divided peak current. Deactivation Tw is the weighted deactivation time. N is the number of HEK293T cells. * = statistically significant after post-hoc correction of p-value for family-wise error; all statistical tests are paired t-tests compared to baseline.

| *CA1 Pyramidal Cells* | | | | | |
|---|---|---|---|---|---|
| | **Rise Time** (ms) | **Peak Amplitude** (pA) | **Tau Weighted** (ms) | **Charge Transfer** (nA*ms) | **Fold Charge Transfer** |
| **Baseline** *(n = 6 cells)* | 5.9 ± 0.5 | -48 ± 4.3 | 82 ± 13 | -4.5 ± 0.6 | 1.0 |
| **10 µM (+)-EU1180-453** *(n = 6 cells)* | 6.2 ± 0.6 | -54 ± 5.7 | 88 ± 11 | -5.4 ± 0.4 | 1.3 ± 0.1 |
| **Baseline** *(n = 6 cells)* | 5.0 ± 0.4 | -79 ± 13 | 58 ± 5.3 | -5.1 ± 0.9 | 1.0 |
| **0.1% DMSO** *(n = 6 cells)* | 5.8 ± 0.4 | -79 ± 5.3 | 63 ± 6.7 | -6.0 ± 0.9 | 1.2 ± 0.1 |
| *CA1 Stratum Radiatum Interneurons* | | | | | |
| | **Rise Time** (ms) | **Peak Amplitude** (pA) | **Tau Weighted** (ms) | **Charge Transfer** (nA*ms) | **Fold Charge Transfer** |
| **Baseline** *(n = 14 cells)* | 7.1 ± 1.3 | -69 ± 11 | 94 ± 15 | -6.2 ± 1.3 | 1.0 |
| **10 µM (+)-EU1180-453** *(n = 14 cells)* | 7.5 ± 1.2 | -93 ± 15* | 120 ± 18* | -10 ± 2.0* | 2.1 ± 0.4 |
| **Baseline** *(n = 5 cells)* | 10 ± 2.5 | -120 ± 13 | 97 ± 16 | -13 ± 2.0 | 1.0 |
| **0.1% DMSO** *(n = 5 cells)* | 9.2 ± 2.2 | -110 ± 9.1 | 110 ± 18 | -12 ± 1.8 | 0.95 ± 0.1 |

**Supplemental Table S4**. Summary data from evoked NMDAR-mediated EPSCs onto CA1 pyramidal cells or *stratum radiatum* interneurons. Baseline and 10 µM (+)-EU1180-453 or baseline and 0.1% DMSO recorded in the same cell. Each neuron was recorded from a different slice. Fold charge transfer is fold change of charge transfer for the last 3 minutes of the drug/vehicle period compared to the last 3 minutes of the baseline period. All data are mean ± SEM; N is the number of neurons. * $p < 0.05$ for a paired t-test compared to baseline, after post-hoc correction of p-value for family-wise error.

| *Wildtype CA1 Pyramidal Cells* | | | | | |
|---|---|---|---|---|---|
| | **Frequency** (Hz) | **Amplitude** (pA) | **Decay Time** (ms) | **Fold Frequency** | **Fold Amplitude** |
| **Baseline** *(n = 11 cells)* | 7.7 ± 1.0 | 23 ± 1.8 | 12 ± 0.5 | 1.0 | 1.0 |
| **10 µM (+)-EU1180-453** *(n = 11 cells)* | 10 ± 1.1* | 24 ± 2.5 | 12 ± 1.0 | 1.4 ± 0.01 | 1.0 ± 0.03 |
| **Baseline** *(n = 11 cells)* | 16 ± 2.4 | 20 ± 0.9 | 14 ± 1.0 | 1.0 | 1.0 |
| **0.1% DMSO** *(n = 11 cells)* | 16 ± 2.0 | 19 ± 1.1 | 15 ± 0.9 | 1.0 ± 0.04 | 1.0 ± 0.05 |
| *Grin2d$^{-/-}$ CA1 Pyramidal Cells* | | | | | |
| | **Frequency** (Hz) | **Amplitude** (pA) | **Decay Time** (ms) | **Fold Frequency** | **Fold Amplitude** |
| **Baseline** *(n = 14 cells)* | 2.1 ± 0.2 | 19 ± 1.7 | -- | 1.0 | 1.0 |
| **10 µM (+)-EU1180-453** *(n = 14 cells)* | 1.7 ± 0.2* | 16 ± 1.5 | -- | 0.8 ± 0.08 | 0.9 ± 0.06 |

**Supplemental Table S5**. Summary of spontaneous inhibitory postsynaptic currents (sIPSCs) onto wildtype or *Grin2d$^{-/-}$* CA1 pyramidal cells in the developing hippocampus. Baseline responses were obtained for 5 minutes then 10 µM (+)-EU1180-453 or vehicle (0.1% DMSO) was applied for 10 minutes. All recordings were finished in 10 µM gabazine and all cells recorded at a holding potential of +10 mV with no other antagonists in the perfusion solution. Each neuron was recorded using a separate slice. Fold frequency is fold change of average frequency for the last 3 minutes of the drug/vehicle period compared to the last 3 minutes of the baseline period. Fold amplitude is fold change of average amplitude for the last 3 minutes of the drug/vehicle period compared to the last 3 minutes of the baseline period. All data are mean ± SEM. N is the number of neurons. * = $p < 0.05$ for a paired t-test compared to baseline, after post-hoc correction of p-value for family-wise error.

| *EPSP/IPSP Recordings* | | | | |
|---|---|---|---|---|
| | **EPSP Amplitude** (mV) | **IPSP Amplitude** (mV) | **EPSP/IPSP** | **Fold EPSP/IPSP** |
| **Baseline** *(n = 8 cells)* | 5.5 ± 1.2 | -2.8 ± 0.4 | 2.0 ± 0.3 | 1.0 |
| **10 μM (+)-EU1180-453** *(n = 8 cells)* | 4.0 ± 1.2* | -4.1 ± 0.6* | 0.9 ± 0.2* | 0.5 ± 0.1 |
| **Baseline** *(n = 4 cells)* | 6.2 ± 0.8 | -5.2 ± 0.6 | 1.3 ± 0.2 | 1.0 |
| **0.1% DMSO** *(n = 4 cells)* | 6.3 ± 1.2 | -6.0 ± 1.0 | 1.2 ± 0.3 | 0.9 ± 0.1 |
| *Carbachol-Induced Gamma Oscillations* | | | | |
| | **Average Power** ($\mu V^2$/Hz, *$10^7$) | **Peak Power** ($\mu V^2$/Hz, *$10^5$) | **Total Power** ($\mu V^2$/Hz, *$10^4$) | **Fold Total Power** |
| **Baseline** *(n = 7 slices)* | 4.8 ± 1.3 | 0.6 ± 0.2 | 0.5 ± 0.2 | 1.0 |
| **20 μM Carbachol** *(n = 7 slices)* | 67 ± 24 | 3.3 ± 1.2 | 6.7 ± 2.4 | 14 ± 3.1 |
| **20 μM Carbachol + 10 μM (+)-EU1180-453** *(n = 7 slices)* | 160 ± 49 | 7.1 ± 2.4 | 16 ± 4.9 | 31 ± 4.0 |
| | **Average γ Power** ($\mu V^2$/Hz, *$10^7$) | **Peak γ Power** ($\mu V^2$/Hz, *$10^5$) | **Total γ Power** ($\mu V^2$/Hz, *$10^4$) | **Fold Total γ Power** |
| **Baseline** *(n = 7 slices)* | 3.1 ± 0.7 | 0.02 ± 0 | 0.2 ± 0 | 1.0 |
| **20 μM Carbachol** *(n = 7 slices)* | 39 ± 15 | 1.7 ± 0.7 | 3.3 ± 1.3 | 14 ± 3.1 |
| **20 μM Carbachol + 10 μM (+)-EU1180-453** *(n = 7 slices)* | 100 ± 34 | 4.4 ± 1.7 | 8.6 ± 2.8 | 31 ± 4.0 |

**Supplemental Table S6**. Summary of excitatory postsynaptic potential (EPSP) and inhibitory postsynaptic potential (IPSP) recordings and carbachol-induced gamma oscillations. For EPSP/IPSP recordings, CA1 pyramidal cells were current-clamped and held at -60 mV. Schaffer collateral stimulation was set 10-20% below threshold to generate a single action potential. Baseline recordings were obtained for 5 minutes and 10 μM (+)-EU1180-453 or 0.1% DMSO was applied for 10 minutes. For carbachol-induced gamma oscillations, field potential recordings from *stratum pyramidale* in CA1 were recorded using an interface-style chamber. Baseline periods were recorded for five minutes, then 20 μM carbachol was applied for 15 minutes, then 20 μM carbachol plus 10 μM (+)-EU1180-453 was applied for another 10 minutes. The last minute of each period was analyzed for power. All data are mean ± SEM. * = $p < 0.05$ for a paired t-test compared to baseline, after post-hoc correction of p-value for family-wise error. γ = gamma band.

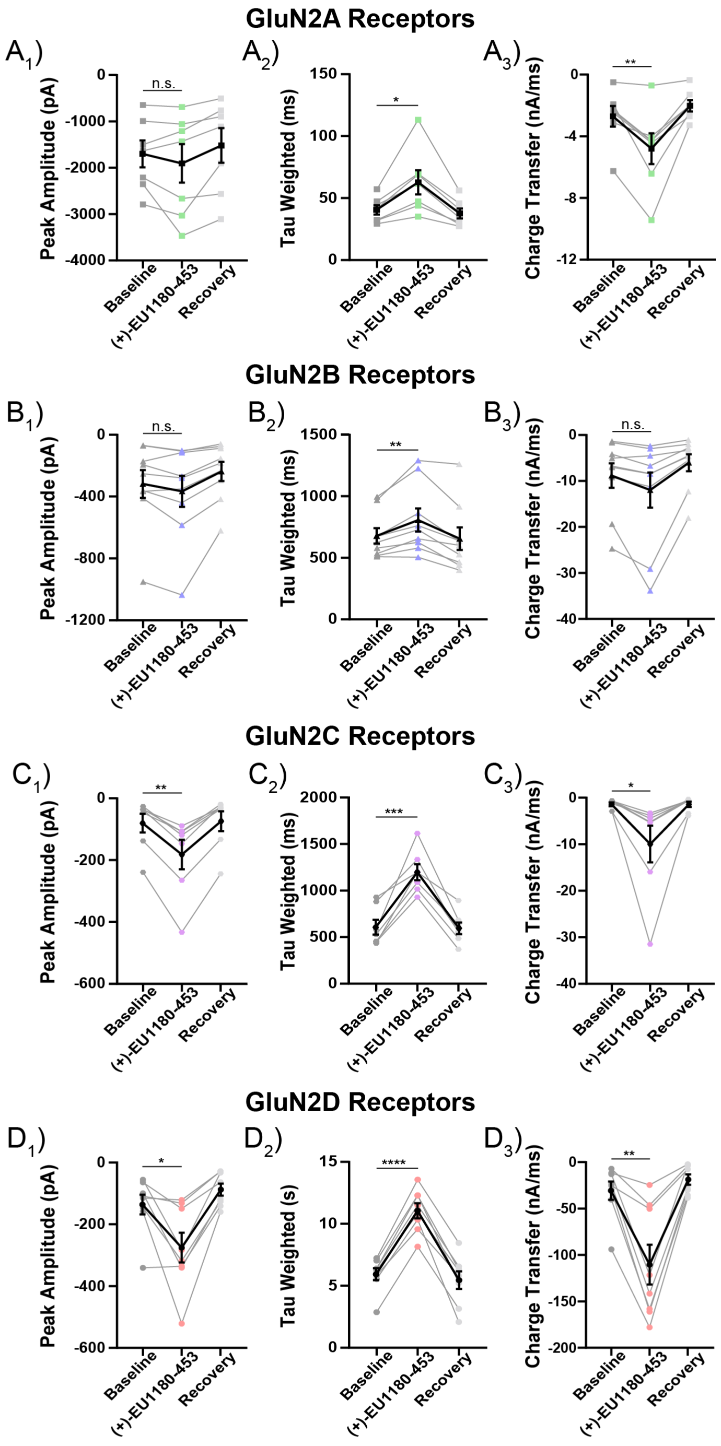

**Supplemental Figure S1.** Summary of response parameters for brief (<10 ms) agonist application. Measured parameters for individual cells (gray) and the mean (black) are shown for response to a maximally effective concentration of coagonists glutamate and glycine in the absence and presence of 10 μM (+)EU1180-453. The peak amplitudes for GluN2A (**A**$_1$) and GluN2B-containing NMDARs (**B**$_1$) were not altered by (+)-EU1180-453, however, the weighted taus describing the deactivation time course for GluN2A- (**A**$_2$) and GluN2B-containing NMDARs (B$_2$) receptors were slightly prolonged by (+)-EU1180-453. **A**$_3$) Charge transfer for GluN2A-containing NMDARs was slightly potentiated whereas that for GluN2B-containign NMDARs was not altered by (+)-EU1180-453. Peak amplitudes for GluN2C- (**C**$_1$) and GluN2D-containing NMDARs (**D**$_1$) were strongly increased by (+)-EU1180-453. Weighted tau values describing the deactivation time course for GluN2C- (**C**$_2$) and GluN2D-containing NMDARs (D$_2$) were prolonged by (+)-EU1180-453. Similarly, charge transfer for both GluN2C- (**C**$_3$) and GluN2D-containing NMDARs (**D**$_3$) was increased by (+)-EU1180-453. See **Supplemental Table S2** for full quantifications. * $p < 0.05$; ** $p < 0.01$; *** $p < 0.001$; **** $p < 0.0001$ for baseline vs (+)EU1180-453, paired t-test.

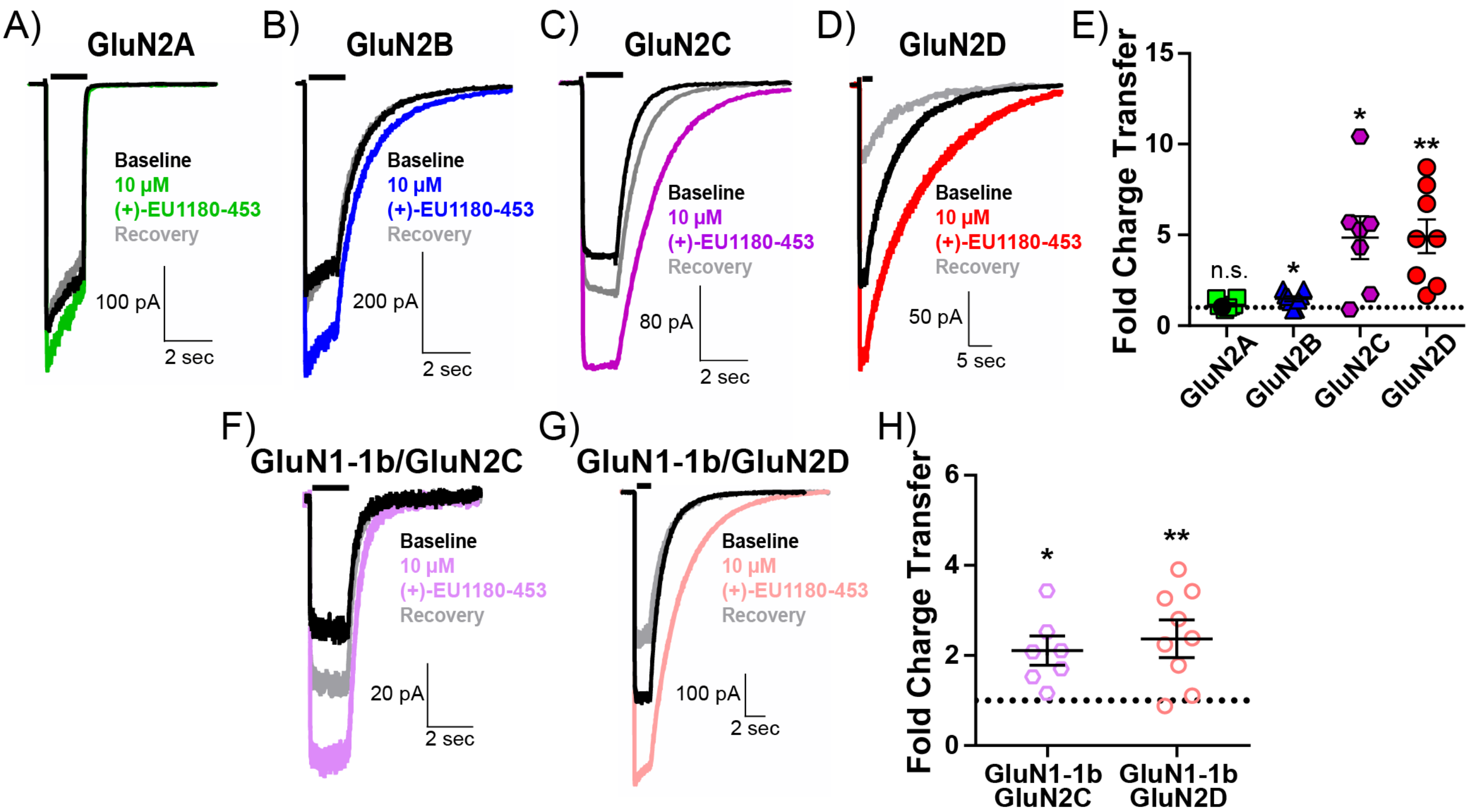

**Supplemental Figure S2.** (+)-EU1180-453 strongly potentiates GluN2C- and GluN2D-containing NMDARs. HEK293T cells were transiently transfected with various subunit combinations of NMDARs and glutamate (100 µM) was rapidly applied for 1.5 sec to establish a baseline response. Cells were then exposed to saturating glutamate (100 µM) and 10 µM (+)-EU1180-453 to obtain a paired modulator response. Recordings finished by removing (+)-EU1180-453 from the solution to demonstrate reversibility of the potentiated response. All recordings were performed in saturating glycine (30 µM) and at a holding potential of -60 mV. Representative responses to baseline (glutamate only, black), glutamate plus 10 µM (+)-EU1180-453 (color), and back to glutamate only (gray) for **A**) GluN2A (n = 8 cells), **B**) GluN2B (n = 9 cells), **C**) GluN2C (n = 7 cells), and **D**) GluN2D (n = 8 cells) diheteromeric NMDARs. **E**) Fold charge transfer (response/baseline) for NMDAR subunit. We observed no change fold charge transfer for GluN2A receptors NMDARs (1.2 ± 0.1 fold change), but minor potentiation of fold charge transfer for GluN2B receptors (1.5 ± 0.1 fold change). Fold change of change transfer for GluN2C receptors and GluN2D receptors was significant and robust, with 4.7 ± 1.4 for GluN2C and 4.9 ± 0.9 for GluN2D. Representative responses to baseline (glutamate only, black), glutamate plus 10 µM (+)-EU1180-453 (color), and back to glutamate only (gray) for **F**) GluN1-1b/GluN2C (n = 7 cells) and **G**) GluN1-1b/GluN2D (n = 9 cells) diheteromeric NMDARs. **H**) Fold change of change transfer for GluN1-1b/GluN2C receptors and GluN1-1b/GluN2D receptors was significant and robust, with 4.8 ± 1.2 for GluN1-1b/GluN2C and 4.7 ± 1.2 for GluN1-1b/GluN2D. All data are mean ± SEM. Statistical significance determined by paired t-test; no statistical tests were performed on any recovery values. See **Supplemental Table S3** for full quantifications. * $p < 0.05$; ** $p < 0.01$; n.s. = not significant.

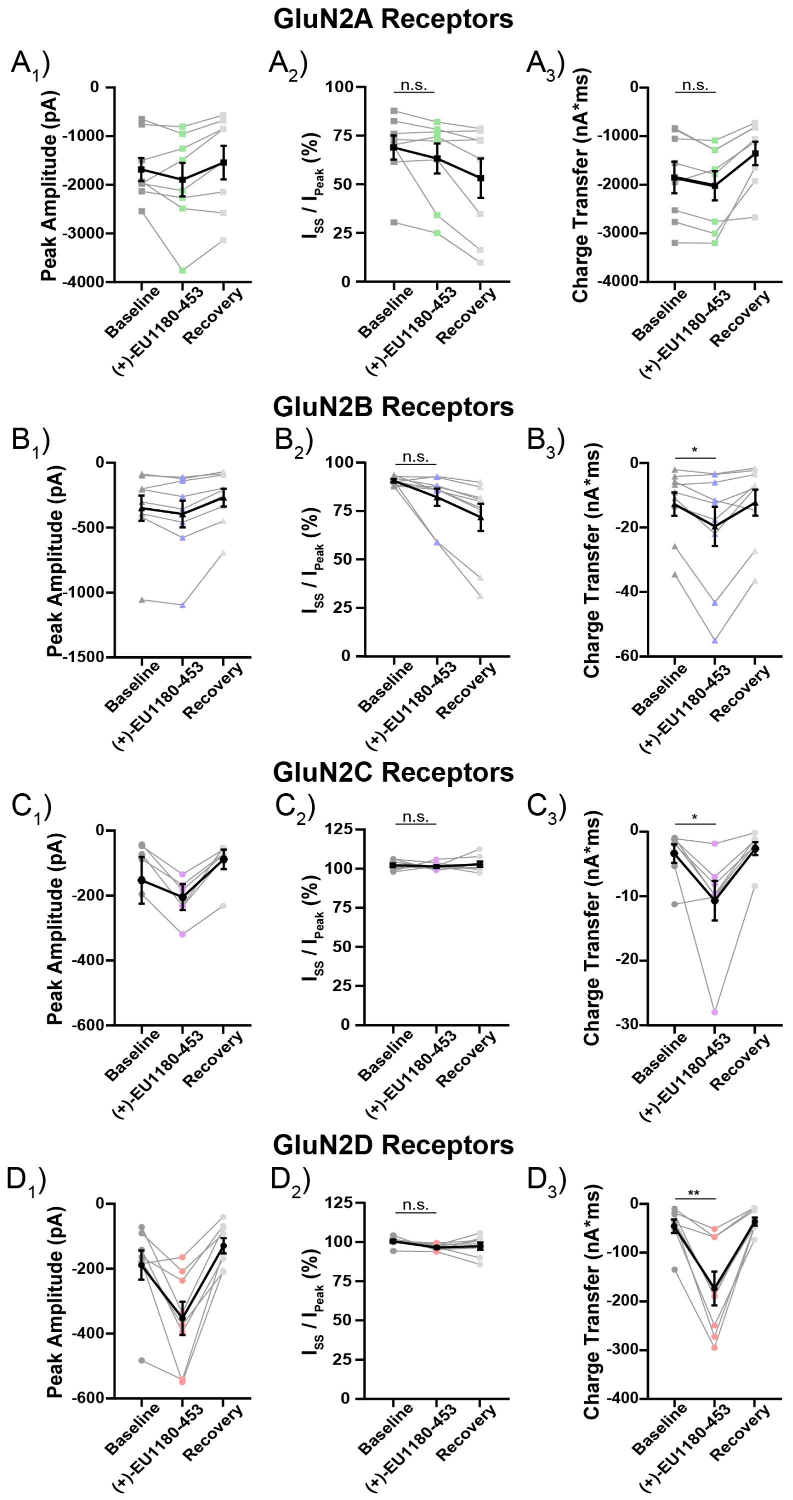

**Supplemental Figure S3.** Summary of response parameters for prolonged <10 ms agonist application Additional data for 1.5 sec HEK cells jumps. **A$_1$, B$_1$, C$_1$, and D$_1$**) Peak amplitude at GluN2A, GluN2B, GluN2C, and GluN2D receptors, respectively. **A$_2$** and **A$_3$**) $I_{SS}/I_{peak}$ and charge transfer, respectively, at GluN2A NMDARs is not altered by (+)-EU1180-453. **B$_2$**) Tau weighted describing the deactivation time course at GluN2B receptors is not altered by (+)-EU1180-453. **B$_3$**) Charge transfer at GluN2B receptors is slightly increased by (+)-EU1180-453. **C$_2$**) $I_{SS}/I_{peak}$ at GluN2C NMDARs is not altered by (+)-EU1180-453. **C$_3$**) Charge transfer at GluN2C receptors is increased by (+)-EU1180-453. **D$_2$**) $I_{SS}/I_{peak}$ ratio used to describe desensitization at GluN2D receptors is not modulated by (+)-EU1180-453. **D$_3$**) Charge transfer at GluN2D receptors is increased by (+)-EU1180-453. All data are mean ± SEM. Statistical significance determined by paired t-test; no statistical tests were performed on peak amplitude or any recovery values. See **Supplemental Table S3** for full quantifications. $I_{SS}$ steady-state current response ~ 100 ms before the end of the 1.5 second glutamate or glutamate plus (+)-EU1180-453 period. $I_{peak}$ peak current obtained immediately after rapid application of into glutamate or glutamate plus (+)-EU1180-453. * $p < 0.05$; ** $p < 0.01$; n.s. not significant.

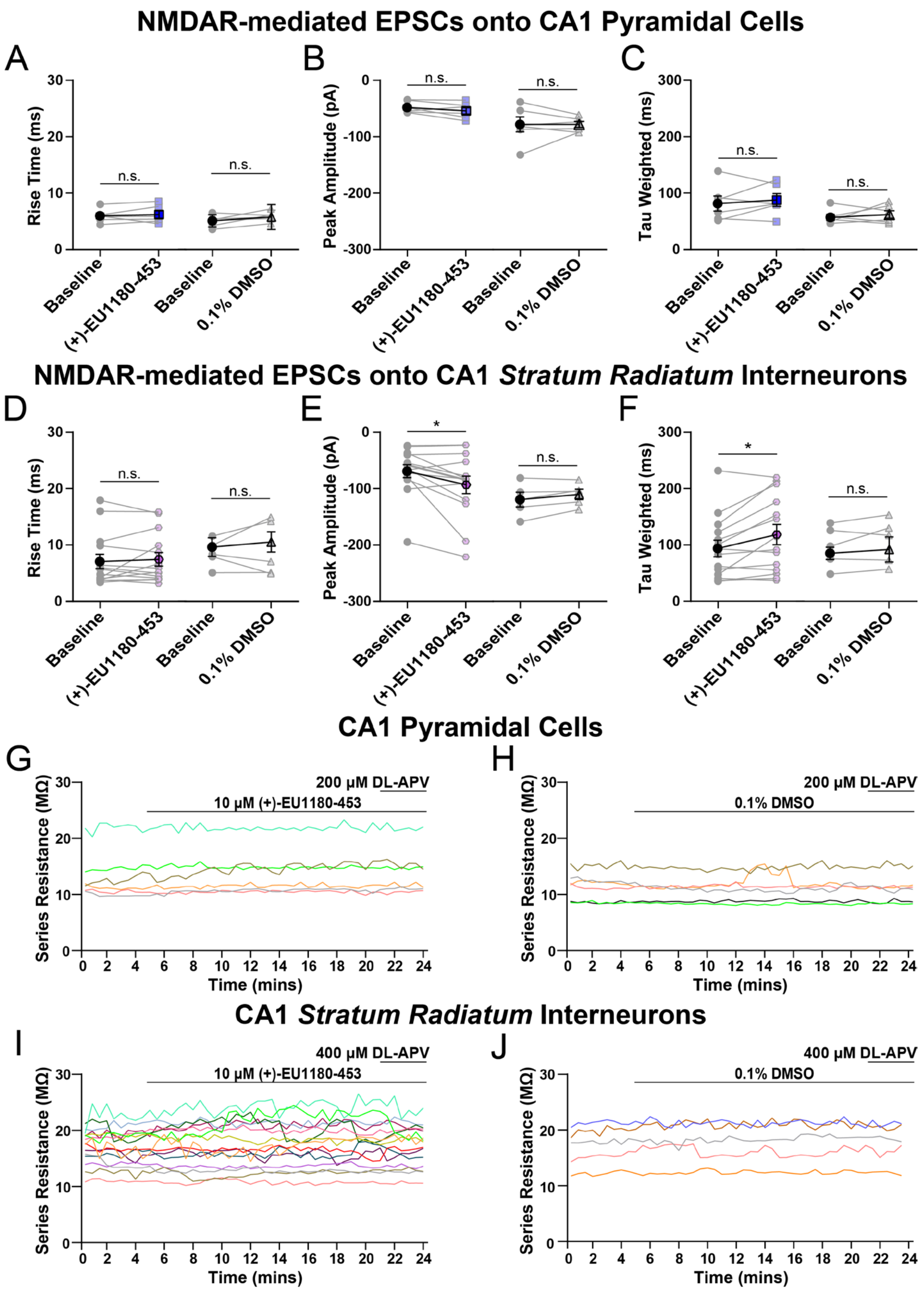

**Supplemental Figure S4**. CA1 *stratum radiatum* GABAergic interneurons, but not CA1 pyramidal cells, are potentiated by (+)-EU1180-453 in developing hippocampus. CA1 pyramidal cells or CA1 *stratum radiatum* GABAergic interneurons were held at -30 mV (0.2 mM $Mg^{2+}$) and NMDAR-mediated excitatory postsynaptic currents (EPSCs) were pharmacologically isolated. Schaffer collaterals were stimulated once every 30 seconds. Baseline responses were recorded for 5 minutes, followed by a 15-minute application period of either 10 μM (+)-EU1180-453 or vehicle (0.1% DMSO). All recordings concluded in 200 μM DL-APV for pyramidal cells or 400 μM DL-APV for interneurons. The last 3 minutes of the baseline period and the drug/vehicle application period were used for statistical analyses. **A**) Rise time, **B**) peak amplitude, and **C**) tau weighted of NMDAR-mediated EPSCs onto CA1 pyramidal cells were all unchanged following application of 10 μM (+)-EU1180-453 (n = 6 cells, blue) or vehicle (0.1% DMSO, n = 6 cells, gray). **D**) Rise times of NMDAR-mediated EPSCs onto CA1 *stratum radiatum* interneurons are unchanged following application 10 μM (+)-EU1180-453 (n = 14 cells, magenta) or vehicle (n = 5 cells, 0.1% DMSO, gray). **E**) Peak amplitude of NMDAR-mediated EPSCs onto CA1 *stratum radiatum* interneurons were increased following application 10 μM (+)-EU1180-453 but not with vehicle. **F**) Weighted tau of NMDAR-mediated EPSCs onto CA1 *stratum radiatum* interneurons was prolonged following application 10 μM (+)-EU1180-453 but not with vehicle. Series resistance was monitored throughout recording and is shown for CA1 pyramidal cells with (+)-EU1180-453 (**G**) or vehicle (**H**) and *stratum radiatum* interneurons with (+)-EU1180-453 (**I**) or vehicle (**J**). Data are mean ± SEM. See **Supplemental Table S4** for full quantifications. * $p < 0.05$ by paired t-test; ns = not significant.

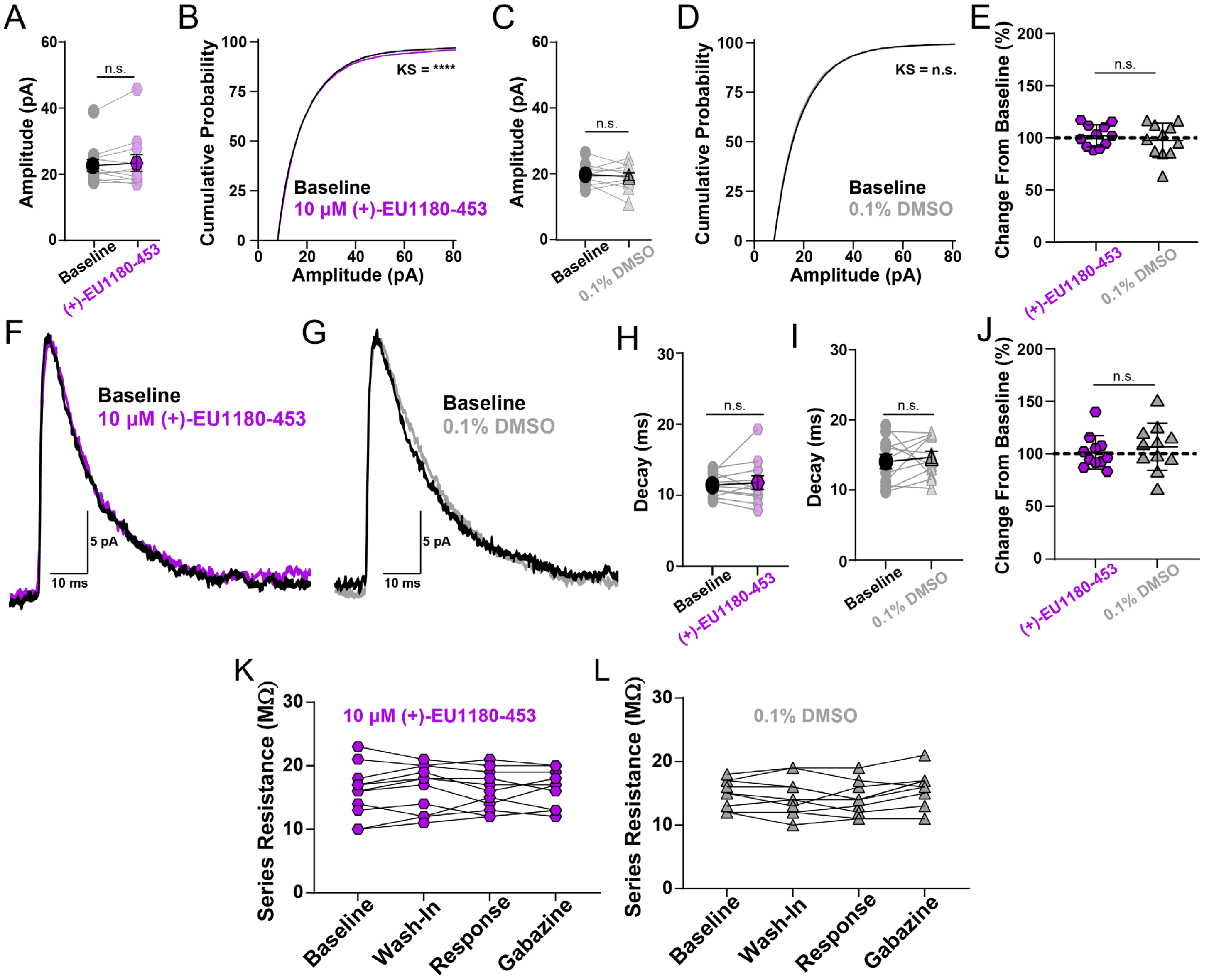

**Supplemental Figure S5.** (+)-EU-1180-453 has minimal effects on spontaneous inhibitory postsynaptic current (sIPSC) amplitude and time course. **A**) Average sIPSC amplitude is unchanged following 10 µM (+)-EU1180-453. **B**) Cumulative probability of event amplitude is significantly shifted to the right after 10 µM (+)-EU1180-453 application (Kolmogorov-Smirnov test, $p<0.0001$). **C**) Average sIPSC amplitude and **D**) cumulative probability of event amplitude are both unchanged in response to vehicle. **E)** Plots of average event amplitude percent change from baseline for 10 µM (+)-EU1180-453 (magenta) and vehicle (gray) no change (unpaired t-test). **F**) Normalized, representative responses of sIPSCs before (black) and after 10 µM (+)-EU1180-453 (magenta) and **G**) before (black) and after vehicle (gray). Average decay time of sIPSCs is unchanged following either **H**) 10 µM (+)-EU1180-453 or **I**) vehicle (0.1% DMSO) application. **J**) Average decay time percent change shows no significant difference between 10 µM (+)-EU1180-453 (magenta) or vehicle (gray). Series resistance was monitored throughout recording of sIPSCs onto CA1 pyramidal cells, and is displayed for **K**) sIPSC recordings with 10 µM (+)-EU1180-453 (magenta) and **L**) sIPSC recordings with vehicle (0.1% DMSO, gray). All data are mean ± SEM. Statistical significance was determined by paired t-test unless otherwise noted. See **Supplemental Table S5** for full quantifications. K.S. Kolmogorov-Smirnov test; **** $p < 0.0001$; n.s. not significant.

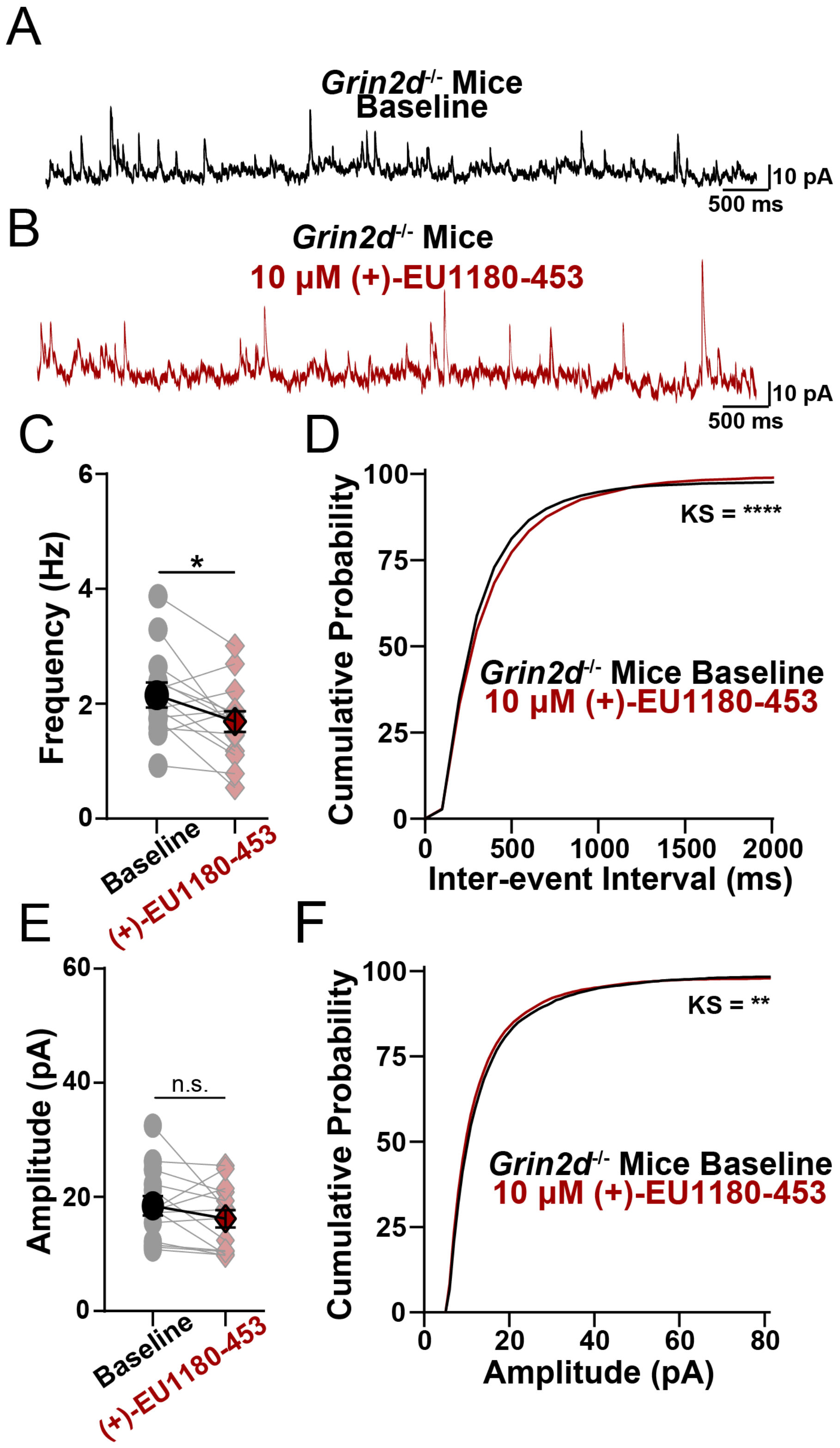

**Supplemental Figure S6**. Effect of sIPSCs after (+)-EU1180-453 application in *Grin2d*$^{-/-}$ mice. Representative current responses of sIPSCs in *Grin2d*$^{-/-}$ mice **A**) before (black) and **B**) after 10-minute wash-in of 10 µM (+)-EU1180-453 (maroon). **C**) There is a significant decrease in average event frequency in *Grin2d*$^{-/-}$ mice in response to (+)-EU1180-453 (paired t-test; $p < 0.05$). **D**) Cumulative probability of event interevent interval is significantly shifted to the right after 10 µM (+)-EU1180-453 application in *Grin2d*$^{-/-}$ mice (Kolmogorov-Smirnov test, $p<0.0001$). **E**) There is no change in the average event amplitude in *Grin2d*$^{-/-}$ mice (paired t-test), however, **F**) the cumulative probability of event amplitude is significantly shifted to the left (Kolmogorov-Smirnov test, $p<0.01$). All data are mean ± SEM. Statistical significance determined by paired t-test unless otherwise noted. See **Supplemental Table S5** for full quantifications. K.S. Kolmogorov-Smirnov test; **** $p < 0.0001$; n.s. not significant.

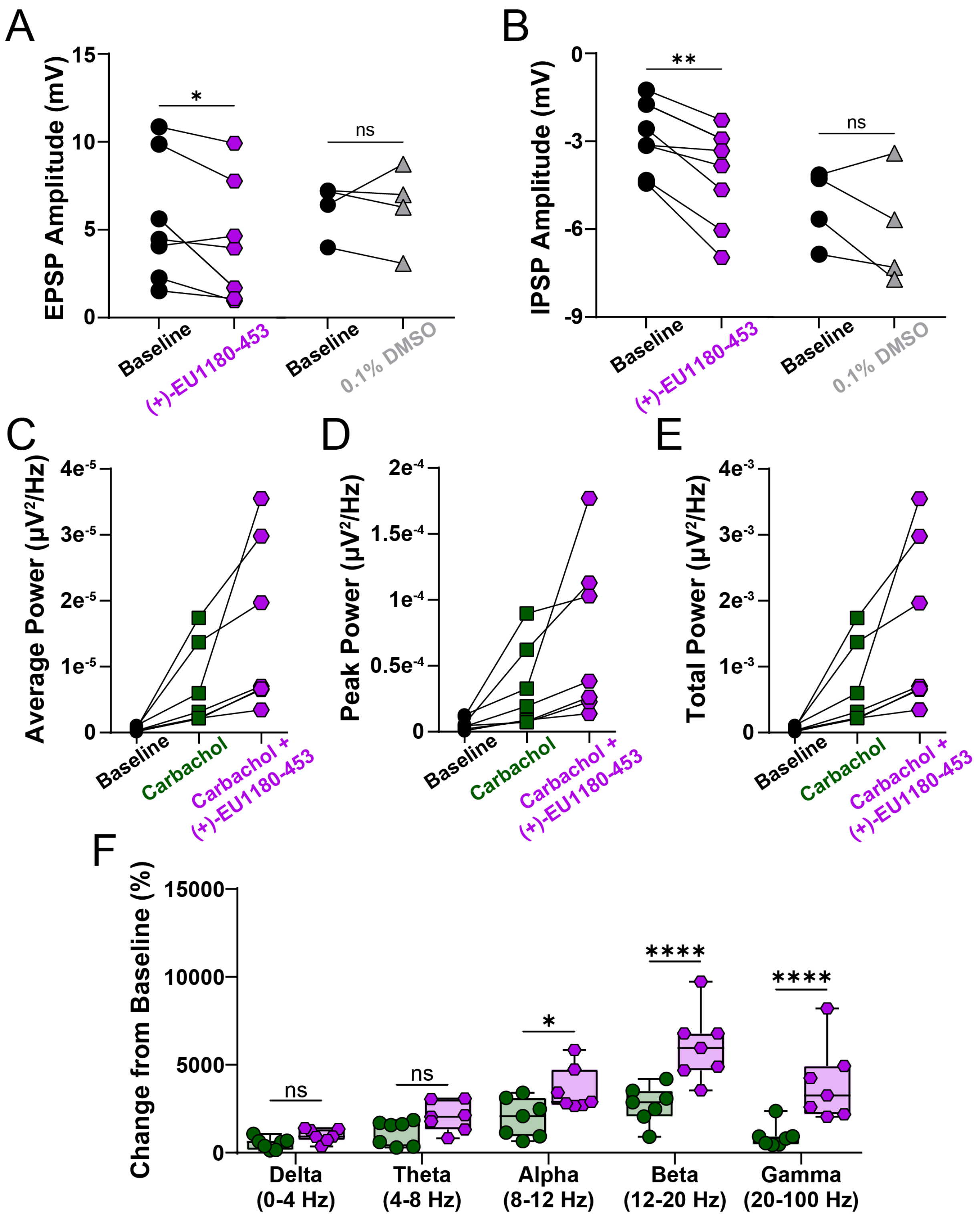

**Supplemental Figure S7.** Additional data on dual excitatory postsynaptic potential (EPSP) and inhibitory postsynaptic potential (IPSP) recordings and carbachol-induced gamma oscillations. **A**) Application of 10 µM (+)-EU1180-453 (magenta) significantly decreases EPSP amplitude (paired t-test), while DMSO leaves EPSP amplitude unchanged. **B**) Application of 10 µM (+)-EU1180-453 (magenta) significantly increases IPSP amplitude (paired t-test), while DMSO leaves IPSP amplitude unchanged. Individual data points from *stratum pyramidale* field potential recordings for baseline (black), 20 µM carbachol (green), and 20 µM carbachol plus 10 µM (+)-EU1180-453 (magenta) showing **C**) average power, **D**), peak power, and **E**) total power. **F**) Percent change in power compared to baseline for 20 µM carbachol (green) and 20 µM carbachol plus 10 µM (+)-EU1180-453 (magenta). Addition of (+)-EU1180-453 significantly enhances power in the alpha, beta, and gamma bands (two-way ANOVA). See **Supplemental Table S6** for full quantifications. All data are mean ± SEM. EPSP = excitatory postsynaptic potential; IPSP = inhibitory postsynaptic potential; * = $p < 0.05$; ** = $p < 0.01$; ns = not significant.
